# Transmission-selective muscle pathology induced by active propagation of mutant huntingtin across the human neuromuscular synapse

**DOI:** 10.1101/2021.07.28.454044

**Authors:** Dinamarca C. Margarita, Laura Colombo, Brykczynska Urszulax, Grimm Amandine, Tousiaki E. Natalia, Fruh Isabelle, Imtiaz Hossain, Gabriel Daniela, Eckert Anne, Müller Matthias, Pecho-Vrieseling Eline

## Abstract

A potential explanation for the spatiotemporal accumulation of pathological lesions in the brain of patients with neurodegenerative protein misfolding diseases (PMDs) is cell-to-cell transmission of aggregation-prone, misfolded proteins. Little is known about central to peripheral transmission and its contribution to pathology. We show that transmission of Huntington’s disease- (HD-) associated mutant HTT exon 1 (mHTTEx1) occurs across the neuromuscular junctions in human iPSC cultures and *in vivo* in *wild-type* mice. We found that transmission is an active and dynamic process, that happens prior to aggregate formation and is regulated by synaptic activity. Furthermore, we find that transmitted mHTTEx1 causes HD-relevant pathology at a molecular and functional level in human muscle cells, even in the presence of ubiquitous expression mHTTEx1. With this work we uncover a casual-link between mHTTEx1 synaptic transmission and pathology, highlighting the therapeutic potential in blocking toxic protein transmission in PMDs.

## Introduction

Neurodegenerative protein misfolding diseases (PMDs) are a group of unrelated illnesses, including Alzheimer’s-(AD), Parkinson’s-(PD), Huntington’s disease (HD), Amyotrophic lateral sclerosis (ALS) and frontotemporal lobar dementia (FTLD). They are all characterized by misfolding and aggregation of a disease-specific protein, cell-type specific vulnerability to degeneration and progressive loss of structure and function of the nervous system. The disease process is already active for years, prior to revealing itself mostly around mid-age with initially discrete neurobehavioral and neuropsychiatric symptoms, which progressively worsen into cognitive impairment^1^. Currently no therapies are available to cure or at least slow down the progression of these devastating illnesses.

It has been suggested that intra brain transmission of the toxic misfolded protein species might be a potential explanation for the spatiotemporal propagation of the pathological lesions through the brain^2^. A casual-link between transcellular spreading of misfolded prion proteins (PrP scrapie or PrPSc) and pathology has been demonstrated in prion diseases^3,4^. For the other neurodegenerative PMDs it has been by now firmly demonstrated that tau (AD), α-synuclein (PD), mutant huntingtin (HD) and tdp-43 (ALS, FTLD) are transmitted between cells and functional connected brain regions (for review see^5-7^). This transmission is accompanied by the appearance of protein aggregates in the acceptor cells. Furthermore, a decline of cognitive and motor behavior has been associated with region-or cell-type specific misfolded protein-expression ^8-12^.

Various cellular mechanism responsible for cell-to-cell transmission of misfolded proteins have been proposed^13^. It has been shown that transneuronal transmission of A-β and tau is enhanced by neuronal activity and synaptic connectivity and that preventing synaptic vesicle release reduces the transmission of mutant huntingtin (mHTT)^14-18^. Together, with the observations that tau, α-synuclein (α-syn), mHTT and tdp-43 are transmitted between functional connected brain regions *in vivo* in mice and drosophila, this strongly suggests a transsynaptic transmission pathway of misfolded proteins^8-11,14,17,19-21^. Synaptic connections are not only present in the central nervous system CNS, but also allow transcellular communication between the CNS and the periphery, as for example the neuromuscular junction (NMJ) between spinal motor neurons and skeletal muscles. mHTT expressed in either the skeletal muscle or brain in Caenorhabditis (C.) elegans has been shown to travel between the CNS and skeletal muscles^22^. Thus, transmission of misfolded proteins could represent a systemic disease pathway affecting not only the CNS, but also contributing to a progressive deterioration of peripheral systems.

Patients with HD suffer from a decline in skeletal muscle function, which progressively worsen with disease course^23,24^. HD is an autosomal dominant disorder that develops with hundred percent penetrance when the number of CAG triplets in the HTT gene exceeds 35 repeats. This repeat is translated into a pathogenic polyglutamine stretch in the exon1 of the HTT protein^25^. Incomplete mRNA splicing of the mHTT results in a toxic exon 1 fragment of the protein, which is highly prone to aggregation and aberrantly translocates to the nucleus where it interferes with transcription^26-29^. Human neuronal cell lines overexpressing the HTT exon 1 (HTTEx1) develop intra-nuclear inclusions and mitochondrial dysfunction^30^. These pathologies are also observed in skeletal muscles of HD patients and animal models, together with skeletal muscle wasting and fatigue^24,31-36^.

Using an isogenic human induced pluripotent stem cell-(hiPSC-) neuromuscular (NM) model combined with high-throughput live-cell imaging, functional analysis and microfluidic systems, we addressed whether mHTTEx1 cell-to-cell transmission can occur across the NMJ and how synaptic density and activity could influence this process. We also examined whether mHTT NM transmission can contribute to skeletal muscle pathology including conditions resembling ubiquitous expression of mHTTEx1. We show that mHTTEx1 is transmitted from neurons to myotubes across the NMJ and that transmission is elevated by increased NMJ density and modulated by neuronal activity. Moreover, our data reveal that transmission occurs independent of mHTTEx1 aggregation, already during NMJs assembly and is enhanced during their functional maturation. Furthermore, our data discloses that mHTTEx1 transmission results in fragmented mitochondria, increased intra-nuclear aggregates and a functional decline of myotube contractibility. Importantly, these pathologies are enhanced or specifically induced by transmission in the presence of cell autonomous mHTTEx1 in myotubes. Finally, we show that mHTTEx1 expressed specifically in the pyramidal neurons in the M1 motor cortex *in vivo* in mice is transmitted to the spinal motor neurons and triceps muscles. Our findings therefore suggest that mHTTEx1 cell-to-cell transmission occurs between the central nervous system and the periphery and might contribute to pathological alterations of the NM system already at early, preclinical stages of the disease. More broadly, our findings also support the notion that cell-type specific vulnerability might be determined by the level of functional synaptic connectivity in combination with trans-synaptic transmission of the misfolded proteins.

## Results

### Characterization of an *in vitro* hiPSC-derived NM co-culture to study transmission of pathogenic HTT

To assess whether mHTT transmission can contribute to skeletal muscle pathology in HD patients, we designed an *in vitro*, isogenic hiPSC-derived NM co-culture system, using two transgenic cell lines, one bearing a doxycycline- (dox)-inducible pro-neuronal transcription factor, *neurogenin 2* (*Ngn2*) transgene^37^ and one bearing a Dox-inducible pro-skeletal muscle transcription factor, *myoblast determination protein 1* (*MyoD*) transgene. We generated four isogenic hiPSC *Ngn2* and one hiPSC *iMyoD* line, isogenic to the *Ngn2* lines, to establish two NM-co-culture systems: **1)** a *Ngn2* line expressing the *exon1* of the HTT gene, with 72 (pathogenic) triplets encoding for glutamine, fused to a *Cre* sequence without the additional nuclear localization signal (*iNgn2;HTTEx1Q72-cre*) and a MyoD hiPSC line with a LoxP-GFP construct (*iMyoD;LoxP-GFP*; Supplemental Fig. 1a). This system can be used for high-throughput, low-resolution, live-cell imaging to follow transmission over weeks (Fig. 1a); **2)** a HTTEx1Q72 fused to a mCherry (*iNgn2;HTTEx1Q72-mCherry;* Supplemental Fig. 1a). The mCherry-tag allows to follow the HTTEx1Q72 transmission to myotubes quantitatively over time. Once a LoxP-GFP myotube turns green, further transmission of HTTEx1Q72 cannot be visualized. In contrast, a change in mCherry labelling reveals the dynamics of this process and allows to correlate the amount of transmitted protein with the pathology. Importantly, using HTTEx1Q72 fused to two different tags we verify that transmission is independent of the tag.

**Figure 1:**
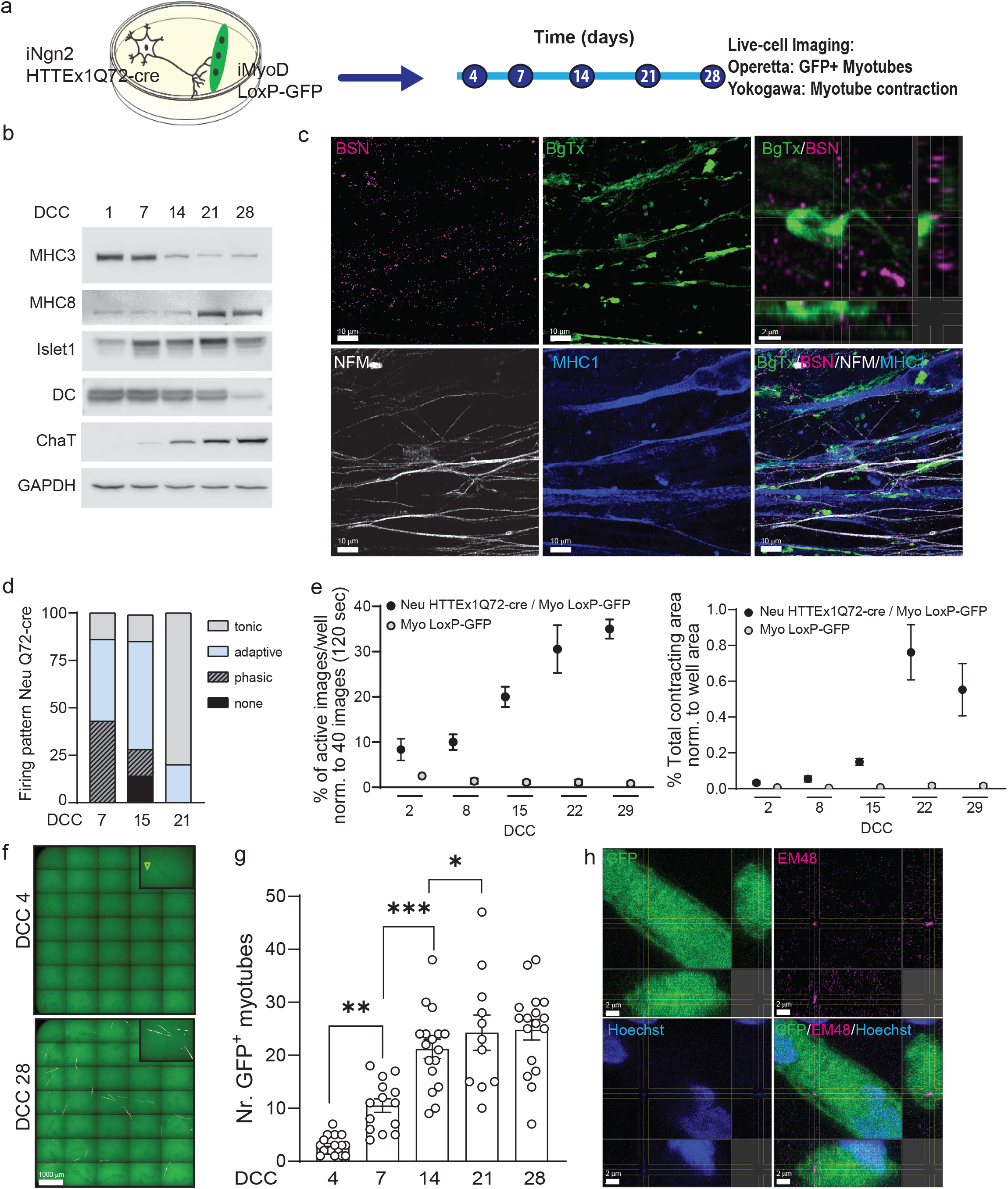
Transmission of HTTEx1Q72 from neurons to muscle cells in hiPSC-derived neuromuscular co-cultures. **a**) Experimental approach to follow in parallel, the development of functional NMJ activity and transmission of HTTEx1Q72-cre from neurons to Myotubes bearing a LoxP-GFP sequence by live-cell, high-throughput imaging. **b**) Representative western blot of developmental markers for myotubes (MHC3 = embryonic myosin, MHC8 = postnatal myosin) and for motor neurons (DC, Islet1 and ChaT) at increasing DCC. **c**) IF images of neuromuscular synapses in Neu HTTEx1Q72-cre / Myo LoxP-GFP co-cultures at DCC21.Top right image: orthogonal view of presynaptic active zone marker BSN in close apposition to the postsynaptic marker BgTx (labels the AChRs on myotubes). NFM labels the axons and MHC1 is a pan-myosin marker. **d**) Distribution of current-induced AP firing patterns of Neu HTTEx1Q72-cre under voltage-clamp of the neurons at -70mV, at DCC 7-21. (n=5-7 neurons per time point) **e**) Percentage of images with myotube contractions (left graph) and total myotube contracting area (right graph) obtained from a single well of a 96-well plate of NM co-culture and muscle only culture, at increasing DCC (n=9 wells/time point in 3 independent cultures (at DCC 2 n=6 from 2 cultures), Two-way Mixed ANOVA (without DCC2): time dependent significant difference between co- and muscle cultures (p=1.21e-10 left panel, p=0.0001 right panel). Post-hoc one-way repeated measures ANOVA: significant increase with time in both parameters for co-culture (p=0.0003 left panel, p=0.0001 right panel) and no significant effect for muscle cultures. **f**) Live-cell fluorescent image from Operetta high-throughput imaging system revealing GFP+ myotubes at DCC 4 (arrowhead) and 28. **g**) Number of GFP+ myotubes obtained by Operetta in Neu HTTEx1Q72-cre / Myo LoxP-GFP co-culture at increasing DCC (n=11-17 wells/time point from at least 4 independent experiments) ***p=0.0008, **p=0.002, *p=0.015 (Linear Mixed Model, Tukey’s correction). **h**) Orthogonal view of an IF image showing an Em48+ HTT aggregate in the cytoplasm of a GFP+ myotube. Hoechst (blue) labels the nuclei. Abbr: **AChRs** = acetylcholine receptors; **BgTx** = a-bungarotoxin; **BSN** = Bassoon; **ChaT** = choline acetyltransferase; **DC** = doublecortin; **DCC** = days of co-culture; **IF** = immunofluorescence; **MHC** = myosin heavy chain; **NFM** = neurofilament M. All averaged data are shown as the mean ± s.e.m.

We assessed transgene expression with western blot (WB) analysis using anti-HTT exon 1 antibody, which revealed that the HTTEx1Q72-cre line expressed the lowest and that we had iPSC clones with different levels of HTTEx1Q72-mCherry expression (Supplemental Fig. 1b). To assess whether the different protein expression levels result in distinct propensity to aggregation we differentiated the hiPSC lines into neurons. At day 1 of differentiation we observed loss of pluripotency with decrease in Oct4 expression (Supplemental Fig. 1c). After 7 and 21 days of differentiation we assessed mHTT aggregates with Em48 antibody which has high affinity for the aggregated form. Aggregation was lowest in HTTEx1Q72-cre line and increased with increasing expression level of the HTTEx1Q72 protein in HTTEx1Q72-mCherry clones (Supplemental Fig. 1d).

To test the cre-lox system, we electroporated *iMyoD;LoxP-GFP* hiPSCs with the *HTTEx1-cre* construct and the *iNgn2;*HTTEx1Q72-cre hiPCSs with a *FloxP-mCherry* plasmid. This resulted in GFP and mCherry expressing cells, resp. In the absence of Cre we never observed GFP expression in the *iMyoD;LoxP-GFP* hiPSCs (n = 3; Supplemental Fig. 1e). Altogether, these analyses demonstrate a successful generation of four hiPSC-lines, which can be used to study cell-to-cell transmission of mHTTEx1.

### Formation of functional NMJs in co-cultures of HTTEx1Q72-cre neurons with LoxP-GFP myotubes

In the next step we established NM co-cultures using a two-step differentiation protocol (Supplemental Fig. 2a). A molecular maturation of the myotubes and neurons in co-cultures was assessed by WB at DCC 1, 7, 14, 21 and 28 using myotube-specific antibodies (myosin heavy chain embryonic and postnatal isoform (MHC3 and MHC8, resp.) and neuronal (doublecortin (DC; neuronal precursor marker) and motor neuron (Islet 1 and choline acetyl transferase (ChaT)), specific antibodies. With increasing co-culture time, we found a decrease in the precursor markers and an increase in the postnatal markers (Fig. 1b). This demonstrates a molecular maturation of the two cell types, hereafter referred to as Neu HTTEx1Q72-cre for the neurons and Myo LoxP-GFP for the myotubes. Immunofluorescence antibody staining (IF) further revealed that NMJs are formed between Neu HTTEx1Q72-cre and Myo LoxP-GFP. We observed close appositions of the neuronal presynaptic active zone marker Bassoon (BSN) and the acetylcholine receptor (AChR) marker α-bungarotoxin (α-BgTx) on myotubes (which represent the postsynaptic structure of the NMJ; Fig. 1c). Patch-clamp recordings from Neu HTTEx1Q72-cre revealed functional maturation of a current-induced action potential firing pattern from a mixed phasic / adaptive at DCC7 to a mainly tonic pattern at DCC 21 (Fig. 1d, Supplemental Fig. 2b). We further demonstrated that myotube contractions disappeared upon addition of the NMJ-activity blocker α-BgTx at DCC21 (Supplemental Fig 2c). These data demonstrate the establishment of functional NMJs between Neu HTTEx1Q72-cre and Myo LoxP-GFP. To gain better insight into the temporal development of these NMJs we followed myotube contractions for 29 days in the same wells of either co-cultures or monocultures of Myo LoxP-GFP. This revealed a temporal increase in both myotube activity and contracting area only in co-cultures (Fig. 1e). In addition, we analyzed the variability of these parameters within one culture well. The variability significantly decreased from DCC 15 onwards in co-cultures, but stayed high in monocultures (Supplemental Fig. 2d), indicative of triggered neuron-induced contractions in co-cultures. Concordantly, DCC15 is also a time of steep increase in myotube contracting area and of neuronal maturation based on a more active AP firing pattern (DCC 14 to 21, tonic firing from 14% to 80%) (Fig. 1d, e). These data together demonstrate the establishment of functional NMJs between Neu HTTEx1Q72-cre and Myo LoxP-GFP in this NM-co-culture system and validates it for addressing the question whether HTTEx1Q72-cre can be transmitted from motor neurons to myotubes across functional NMJs.

### Neuromuscular transmission of HTTEx1Q72-cre occurs with time and in the absence of aggregates in the neurons

To assess HTTEx1Q72-cre transmission, we performed high-throughput live-cell fluorescent imaging from the same wells from DCC 4 to 28. At DCC 4 first GFP+ myotubes appeared and their number increased with co-culture time until day 21, after that the number stayed stable (Fig. 1f, g). This timing correlated with establishment of functional NMJs (Fig. 1d, e). Based on Em48 staining at DCC 7 and 21 we did not observe aggregated form of HTT in HTTEx1Q72-cre neurons (Supplemental Fig. 1d). When we stained the co-culture at DDC 28 we detected presence of few Em48 positive aggregates selectively in GFP+ myotubes (Fig. 1h). Transmission thus likely occurs in a non-aggregated form and aggregation takes place in the myotubes.

To prove that HTTEx1Q72-cre NM transmission requires direct cell-cell contact and is not transferred via the culture media, we placed a two-chamber cell culture insert w/o bottom in one dish to allow physical separation of Neu HTTEx1Q72-cre from Myo LoxP-GFP, while the medium was shared. After attachment of the cells, the inserts were removed (DCC1). The surface between the inserts was not coated to prevent the movement of the cells and extension of the axons to the myotubes. In these co-cultures we never observed GFP+ myotubes (Supplemental Fig. 3a, b).

Taken together, the NM co-culture system allows to follow pathogenic HTTEx1 cell-to-cell transmission over weeks with high-throughput low-resolution live-cell imaging. Furthermore, with the expression of GFP in Myo LoxP-GFP we demonstrated that HTTEx1Q72-cre is transmitted from neurons to muscles and can enter the cytosol and the nucleus of the myotubes.

### NM-co-cultures in microfluidic devices reveal HTTEx1Q72 transmission across the NMJs

To demonstrate that HTTEx1Q72 is transmitted across the NMJ and assess whether these structures play a role in determining the efficiency of pathogenic HTT transmission we established NM co-cultures in microfluidic devices (MFDs). These devices allow to co-culture two cell populations in two isolated compartments, connected with microgrooves through which axons can grow and reach the other compartment, allowing them to form connections with myotubes (Fig. 2a, upper panel). To quantify the HTTEx1Q72 NM transmission we used here the co-cultures of Neu HTTEx1Q72-mCherry clone#75 with Myo LoxP-GFP (we kept using this myotube line, but to prevent confusion we will refer to it as Myo when we used it in co-culture with Neu HTTEx1Q72-mCherry). The mCherry labeling of the HTTEx1Q72-mCherry expressing neurons showed that these projected their axons from the presynaptic neuronal compartment to the postsynaptic myotube compartment (Fig. 2a, lower panel). In the myotube compartment NMJs were established, as visualized with IF staining’s of BSN and AChR appositions at DCC21 (Fig. 2b). AChR clusters on the surface of myotubes, can be classified based on their shapes^38^. We performed a detailed shape analysis of these clusters at DCC21 and classified them into four categories: small&elongated, small&round, big&elongated, big&round (Supplemetal Fig. 4a-c). When we compared the clusters of myotubes in mono versus co-cultures we found that the clusters of small&round type were the majority in both cultures but, there was a significant increase in the density of both types of big clusters in the co-cultures (Supplemental Fig. 4d, e). The big-cluster types are thus likely those constituting the NMJs. Supporting this notion, we found that big cluster types were over-represented among those associated with the presynaptic marker BSN, compared to the clusters w/o BSN (Fig. 2c). With these analyses we demonstrate the presence of structural NMJs in the co-cultures grown in MFDs. To judge whether Neu HTTEx1Q72-mCherry clone#75 neurons are able to potentially trigger myotube contractions we performed patch-clamp recordings and found that these cells develop from DCC 7 to 21 a more active firing pattern of current-induced action potential from mainly phasic at DCC 7 to mainly adaptive at DCC 21 (Fig. 2d).

**Figure 2:**
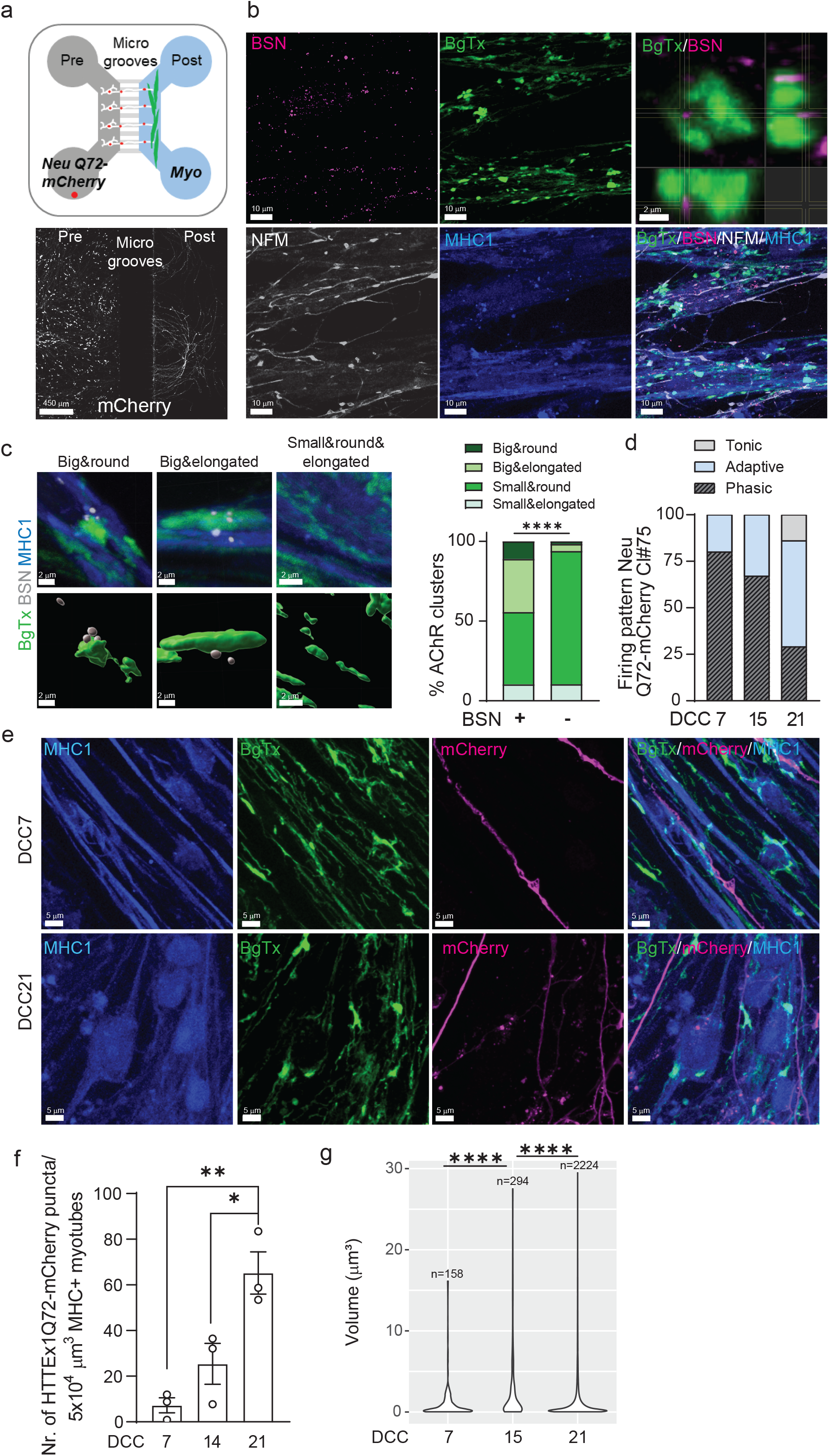
Transmission of HTTEx1Q72 across the NMJ in Neu HTTExQ72-mCherry cl#75 / Myo co-cultures in MFDs. **a, upper panel**) Schematic of MFD depicting the co-culture setting: Neu HTTEx1Q72-mCherry (white with red dots) in presynaptic chamber and Myo (green) in postsynaptic chamber. Neurons extend their axons through the microgrooves to the myotubes. **a, lower panel**) IF of mCherry (white) in MFD visualizing Neu HTTEx1Q72-mCherry in presynaptic chamber and extension of their axons to the postsynaptic compartment. **b**) IF images of neuromuscular synapses in Neu HTTEx1Q72-mCherry cl#75/ Myo co-cultures in MFD at DCC21. Top right image: orthogonal view of close apposition between BSN and BgTx. **c, left)** IF Images of four different types of AChR clusters identified on Neu-Myo co-cultures (upper panels) and corresponding surfaces created in Imaris: AchR clusters visualized by BgTx (green), BSN (white) (lower panels) **c, right**) Distribution of AChR cluster types in percentage, found on Myo when associated with (+) or w/o (-) BSN. (n=1040 for (+); n=11745 for (-)) ****p< 2.2×10^−16^ (χ^2^ test) **d**) Distribution of current-induced AP firing patterns of Neu HTTEx1Q72-mCherry under voltage-clamp of the neurons at -70mV, at DCC 7, 15 and 21. (n=5-7 neurons per time point) **e**) IF of Myo compartment showing the presence of HTTEx1Q72-mCherry+ axons at DCC 7 and axons and puncta at DCC21 in MHC1+ Myo. **f)** Number of HTTEx1Q72-mCherry puncta in Myo at DCC 7 – 21. (n=3 independent co-cultures/time point, data point: mean of 15 images) *p=0.02, **p=0.004 (One-way ANOVA, Tukey’s correction for multiple comparisons) **g**) Violin plot of the volume of individual HTTEx1Q72-mCherry puncta localized in Myo at DCC 7 – 21. n is number of puncta found in Myo across all images at a given time point. ****p ≤5 1×10^−9^(Kruskal-Wallis test, BH correction). Abbr: **AChRs** = acetylcholine receptors; **BgTx** = a-bungarotoxin (labels AChRs); **BSN** = Bassoon; **DCC** = days of co-culture; **IF** = immunofluorescence; **MHC** = myosin heavy chain, **MFD** = microfluidic device. All averaged data are shown as the mean ± s.e.m.

Next, using IF labeling, we revealed that HTTEx1Q72-mCherry protein is present in the myotubes in the postsynaptic compartment, indicating transmission from the neurons (Fig. 2e). Similar as for the HTTEx1Q72-cre, we observed a continuous transmission of HTTEx1Q72-mCherry to Myo, increasing from DCC 7 to DCC 21 quantified as number of HTTEx1Q72-mCherry puncta in the myotubes (Fig. 2f). We analyzed the volume of HTTEx1Q72-mCherry puncta over time and found that there is a dynamic change of a size distribution (Fig. 2g). At DCC 14 we saw appearance of larger aggregates with sizes above 15 µm^3^ and loss of major contribution of smaller assemblies comparing to DCC7. At DDC 21 there was again a major contribution of small assemblies with sizes below 1 µm^3^, together with a persistent presence of bigger aggregates. This suggests that new molecules arrive into the muscle as small assemblies and that aggregation occurs over time. Together, with these experiments we provide evidence that HTTEx1Q72 is transmitted across the NMJ, most probably in a form of small protein complexes.

### The load of HTTEx1Q72-mCherry correlates with increasing NM connections

One of the presymptomatic pathologies in HD patients is a loss of functional neuronal connectivity, that first arises in the cortico-striatal pathway and then progresses to cortical and other subcortical brain regions^39-43^. Interestingly, the most vulnerable brain regions form a selective network with higher connectivity than other brain regions^44^ - a so called ‘rich club’. To assess whether a higher density of NMJ connections leads to more HTTEx1Q72-mCherry puncta in myotubes, we divided the postsynaptic compartment into 3 bins, starting with bin 1 closest to the microgrooves. The area of neuronal processes was highest in bin 1 and decreased towards bin 3 (Fig. 3a, b, Supplemental Fig. 5a). Similarly, we found that the number of HTTEx1Q72-mCherry puncta in myotubes at DCC21 was highest in bin 1 and steeply decreased towards bin 3 (Fig. 3a, c). Interestingly, when we compared the distribution in bins at all time points we observed, a delayed increase in the number of puncta in bin 2 compared to bin 1 (Supplemental Fig 5b). Axons will arrive slightly later in bin 2 than bin 1, followed by delayed formation of NMJs, explaining the temporal delay in HTTEx1Q72-mCherry accumulation in myotubes. Further validating our assumption that HTTEx1Q72-mCherry proteins reach the myotubes via the NMJs, we found a positive correlation between the density of NMJs (BSN-BgTx complexes) and the number of HTTEx1Q72-mCherry puncta in myotubes (Fig. 3d, e).

**Figure 3:**
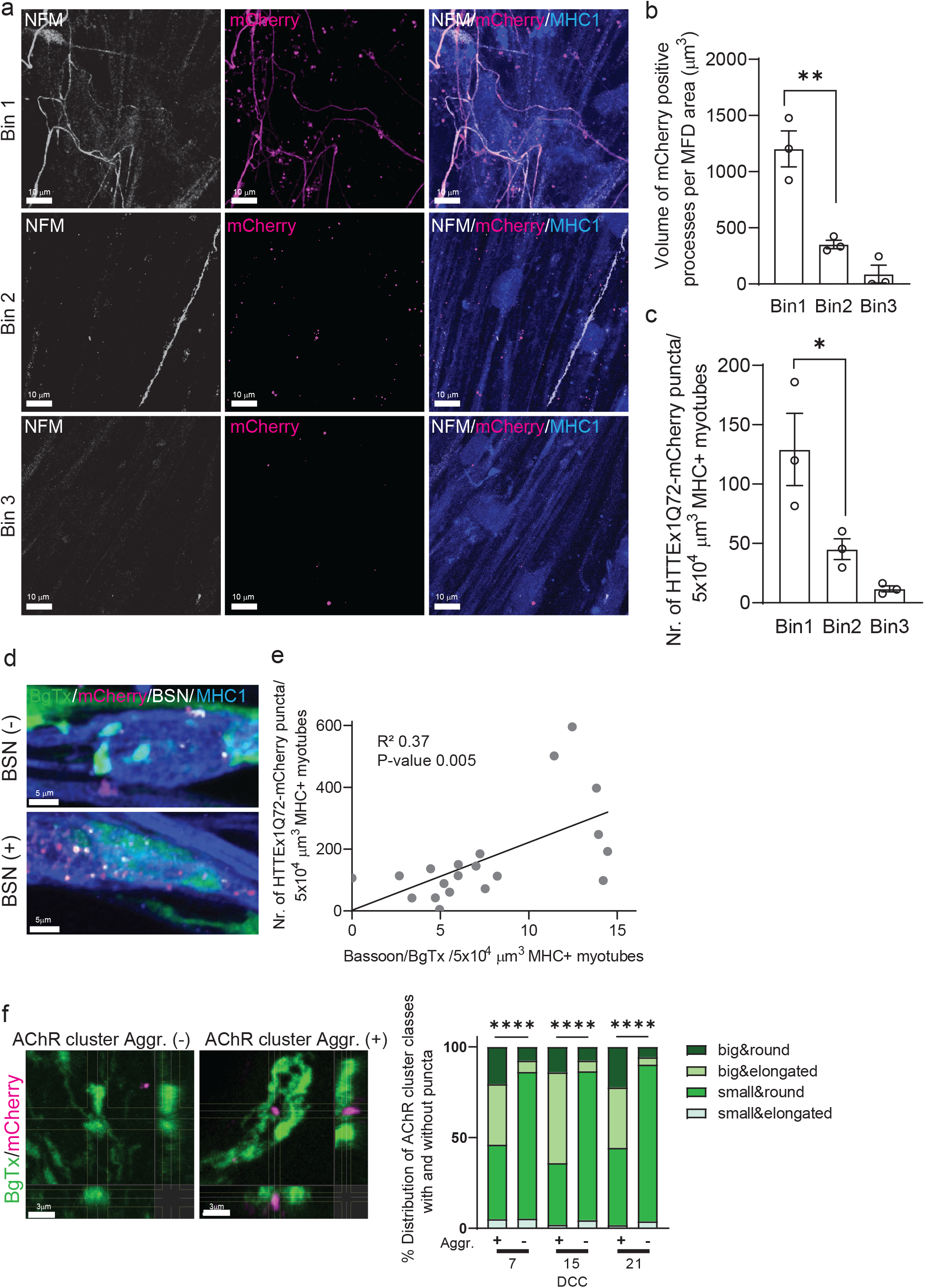
Neuromuscular transmission of HTTExQ72 is enhanced by increasing NMJ density. **a**) IF image showing NFM+ axons and HTTEx1Q72-mCherry in bin 1, bin 2 and bin 3 of Myo compartment in co-cultures with Neu HTTEx1Q72-mCherry cl#75 at DCC21. **b**) Total volume of mCherry positive neuronal processes crossing to Myo compartment normalized to MFD area in each bin. (n=3 independent co-cultures, data point: mean of 5 images) **p=0.003 **c**) Number of HTTEx1Q72-mCherry aggregates inside myotubes in each bin (n=3 independent co-cultures, data point: mean of 5 images) *p=0.04 (One-way ANOVA, Tukey’s correction). **d**) F of Neu Q72-mCherry Myo co-culture showing mHTTEx1 in regions with low (top) or high (bottom) number of BSN-BgTx appositions. **e**) Correlation between number of NMJs (defined as BSN-BgTx appositions on MHC1+ Myo) and number of HTTEx1Q72-mCherry puncta (n=20 images from 3 MFDs, simple linear regression). **f, left panels**) IF of AChR clusters in absence (-) (> 0.05 μm) or close proximity (+) (within 0.05 μm) of HTTEx1Q72-mCherry puncta. **f, right panel**) Distribution of AChR cluster types when associated (within 0.05 μm) with (+) or w/o (-) HTTEx1Q72-mCherry at DCC 7 – 21. (DDC7: n=39 (+); n=10183 (-); DDC15: n=50 (+); n=5161 (-); DDC21: n=277 (+); n=10263 (-)) ****p<0.0001 (Fisher’s Exact Test for DCC7 and 15, χ^2^ test for DCC21). Abbr: **AChRs** = acetylcholine receptors; **BgTx** = a-bungarotoxin (labels AChRs); **BSN** = Bassoon; **DCC** = days of co-culture; **IF** = immunofluorescence; **MHC** = myosin heavy chain; **NFM** = neurofilament M, **MFD** = microfluidic device. All averaged data are shown as the mean ± s.e.m.

Previously we showed that mHTT is transmitted from mouse cells in HD-derived mouse organotypic brain slices (OTBS) to human stem cell-derived neurons (h-neurons). During time of transmission mHTT co-localized with the presynaptic marker synaptophysin and post-synaptic density protein-95 (PSD-95) in neurons^17^. In the NM-co-cultures we also observed around 20% of HTTEx1Q72-mCherry puncta to be associated with the postsynaptic AChRs (Supplemental Fig. 4b). Interestingly, around 60% of the AChR clusters associated with HTTEx1Q72-mCherry puncta were of the big-type, while among those w/o HTTEx1Q72-mCherry only around 10 % were big (Fig. 3f). The big clusters are likely representing those incorporated in the NMJs, since this type increased in the presence of neurons and also in association with the presynaptic marker BSN (Supplemental Fig. 4e, Fig. 2c). Summarizing, we observe a positive correlation of NM-connectivity with HTTEx1Q72-mCherry load in myotubes and preferential association of HTTEx1Q72-mCherry with NMJ-forming AChR clusters.

### Modulating neuronal activity alters NM transmission of mHTTEx1

Neuron-to-neuron transmission of mHTT in the OTBS – h-neuron co-cultures and also *in vivo* in drosophila has been shown to be vastly blocked by preventing SNARE-dependent fusion of synaptic vesicles to the presynaptic membrane and subsequent release of its content ^14,17^. Therefore, we applied the SNARE-cleaving Tetanus neurotoxin (TeNT)^45^ at DCC 10 in HTTEx1Q72-mCherry clone#75 Myo co-cultures. Similar to neuron-to-neuron transmission we observed a significant decrease in HTTEx1Q72-mCherry NM transmission, measured by number of mCherry foci within the myotubes at DCC 21 (Fig. 4a,b). Interestingly, the proportions of AChR cluster types were not affected (Fig.4c), confirming that the observed effect is due to the blocking of the presynaptic neuronal terminals and not due to a re-organization of postsynaptic structures. We hypothesized that synaptic transmission of mHTTEx1 can be regarded as a clearance mechanism by which cells get rid of toxic protein species. We therefore addressed if blocking of this rescue process by the TeNT treatment increased pathological consequences in neurons. Previously, we showed that mHTTEx1 transmitted from mouse cells to human stem cell-derived neurons, first appeared as cytoplasmic aggregates and with time aggregates appeared in the nucleus. Nuclear aggregation correlated with the time when pathological changes occurred in the human neurons^17^. Indeed, in our co-culture system TeNT treatment resulted in increase of the number and size of neuronal intra-nuclear assemblies (Fig. 4d,e).

**Figure 4:**
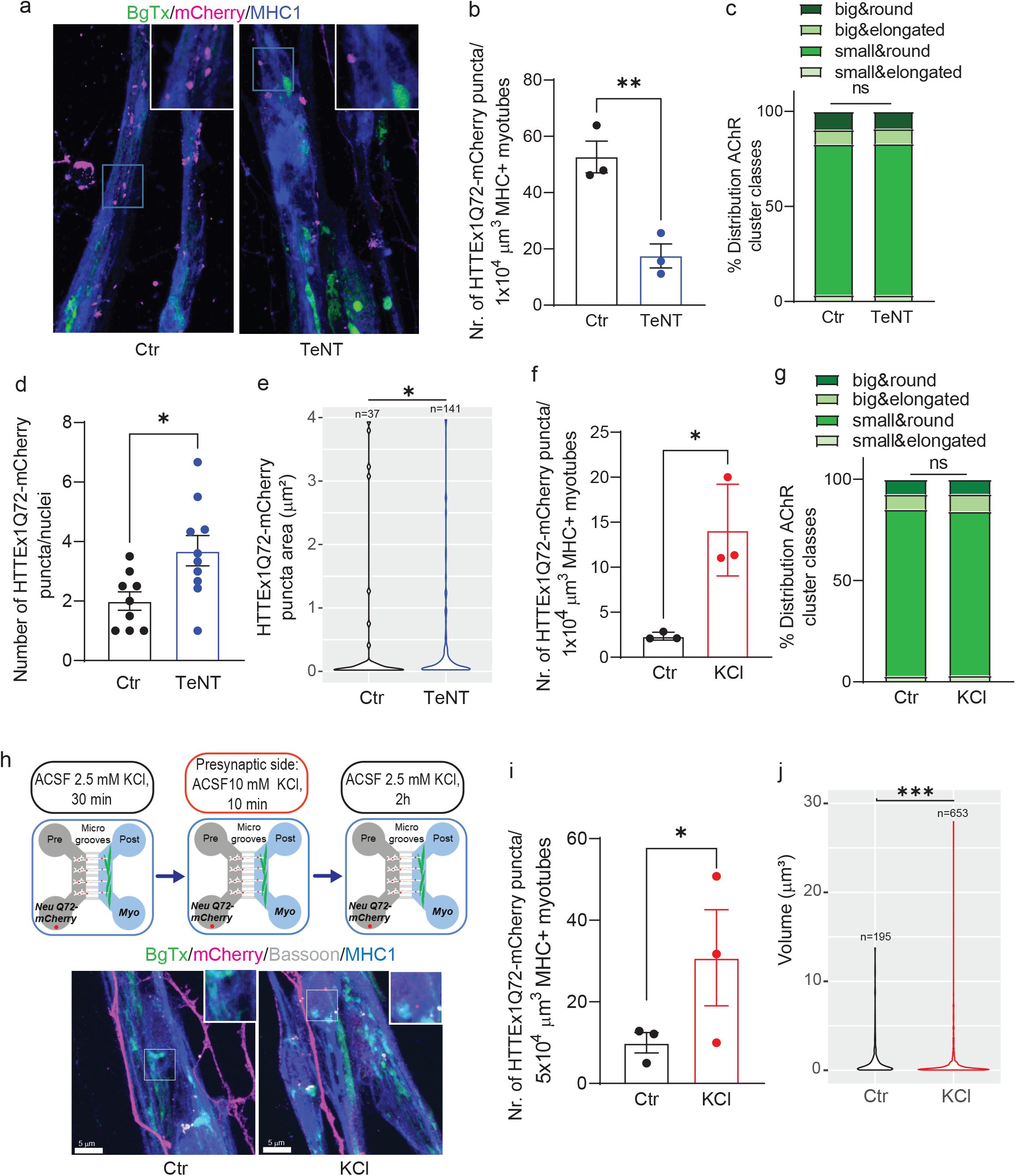
Synaptic activity modulates transmission of HTTEx1Q72 from neurons to myotubes. **a**) IF of Neu HTTEx1Q72-mCherry cl#75/ Myo co-cultures at DCC21: control co-cultures and exposed to 2 μg/ml Tetanus neurotoxin (TeNT) from DCC10. **b**) Number of HTTEx1Q72-mCherry puncta in MHC1+ Myo in control and 2 μg/ml TeNT treated co-cultures (n=3, one data point corresponds to one independent co-culture and is a mean of 5 images, **p=0.0076, Student’s t-test). **c**) Distribution of AChR cluster types, found on Myo in control and TeNT treated co-cultures (n=1304 for (Ctr); n=882 for (TeNT), ns, χ^2^ test). **d**) Number of HTTEx1Q72-mCherry puncta per neuronal nucleus from control or TeNT treated co-culture at DCC21 (n=10 neuronal nuclei/condition, *p=0.013, Student’s t-test). **e**) Violin plot of the area of HTTEx1Q72-mCherry puncta in neuronal nuclei in control and TeNT treated co-cultures, (n indicate the total number of puncta analyzed, *p=0.014, Wilcoxon rank sum test). **f**) Number of HTTEx1Q72-mCherry aggregates in MHC1+ Myo in control and 10 mM KCl treated co-cultures (n=3, one data point corresponds to one independent culture and is a mean of 5 images, *p=0.016, Student’s t-test) **g**) Distribution of AChR cluster types, found on Myo in control conditions or 10 mM KCl treated co-culture (n=1091 for (Ctr); n=1377 for (KCl), ns, χ^2^ test). **h, upper panel**) Schematic outline of experimental approach in MFDs. ACSF with 2.5 mM KCl was added to both the neuronal and myotube compartment for 30 minutes (left schematic). ACSF with 10 mM KCl was added only to the neuronal compartment for 10 minutes (middle schematic). ACSF with 10 mM KCl was changed back to ACSF with 2.5 mM KCl (right schematic). Cultures were fixed after 2 hours. **h, lower panel**) IF images depicting HTTEx1Q72-mCherry puncta in MHC+ Myo in cultures exposed to 2.5 mM KCl (Ctrl) and 10 mM KCl (KCl). The inserts correspond to zoom of a representative region. **i**) Number of HTTEx1Q72-mCherry puncta in MHC1+ Myo in control and 10 mM KCl treated co-cultures. (n=3, one data point corresponds to one independent co-culture and is a mean of 10 images, *p=0.043, Student’s paired t-test). **j**) Violin plot of the volume of HTTEx1Q72-mCherry puncta in control and 10 mM KCl treated co-cultures (n indicate the total number of puncta analyzed, ***p=0.0008, Wilcoxon rank sum test). Abbr: **AChRs** = acetylcholine receptors; **BgTx** = a-bungarotoxin (labels AChRs); **BSN** = Bassoon; **DCC** = days of co-culture; **IF** = immunofluorescence; **MHC** = myosin heavy chain, **MFD** = microfluidic device. All averaged data are shown as the mean ± s.e.m.

As inhibition of synaptic vesicle release reduced HTTEx1Q72-mCherry transmission, we asked if opposite effect can be obtained by depolarization of neurons, triggering increased AP firing and synaptic vesicle release^46^. We exposed the HTTEx1Q72-mCherry clone#72 Myo co-culture at DCC 21 for 10 minutes to 10mM KCl, followed by 2 hours in 2.5mM KCl. We chose to use the clone#72 for this experiment, because of its lower expression of HTTEX1Q72-mCherry, which provides a larger range for increase in transmission without a risk of system saturation. Upon exposure to 10mM KCl we observed more HTTEx1Q72-mCherry puncta in myotubes compared to control co-cultures exposed to 2.5mM KCl (Fig.4f). This increase was not associated with a re-organization of the postsynaptic structures, as AChR proportions did not change after KCl treatment (Fig 4g). To further exclude influence of muscle depolarization on this effect we repeated the experiment in MFDs and applied the 10mM KCl only to the pre-synaptic side (Fig.4h upper panel). We again observed increase in HTTEx1Q72-mCherry puncta in myotubes upon treatment (Fig.4h lower panel, i). When we compared the volume of these HTTEx1 Q72-mCherry puncta, we observed larger contribution of small assemblies in KCl treated compared to non-treated co-cultures, supporting transmission in a form of small protein complexes (Fig. 4j).

The above results of two opposite synaptic manipulations demonstrate that synaptic activity regulates HTTEx1Q72-mCherry transmission and that the pathway of transmission is coupled with synaptic vesicle release.

### HTTEx1Q72-mCherry aggregates accumulate at the myosin surface

Our results so far demonstrate a trans-NM pathway of HTTEx1Q72 transmission. To reveal a potential pathology triggered by transmitted HTTEx1Q72, we assessed the intracellular localization of this protein in the myotubes from DCC 7 to 21. In particular, we analyzed the localization of the HTTEx1Q72-mCherry puncta at the cellular surface. We used the surface function of the Imaris software (Oxford Instruments) to define the surface of myotubes based on MHC1 staining. By visual inspection of images, we observed a striking localization of the puncta to and partially passing through the MHC1+ myotube surface (Fig. 5a, b). Based on a quantification, at DCC 7 half of the puncta localized at the myotube surface (at the surface: 0 – 0.05 µm to myotube surface) and half inside the myotube (inside: > 0.05 µm to myotube surface) and were mostly small (majority below 4 µm^3^) (Fig. 5c, d). At DCC 15 even more - 79% of HTTEx1Q72-mCherry puncta accumulated at the surface, at DCC 21 this shifted back to 52% (Fig. 5c, d). Furthermore, as we found before, we observed growing number of HTTEx1Q72-mCherry puncta with larger volume (Fig. 5c). It has been shown that mHTT has high affinity for bioengineered lipid membranes and that insertion of these proteins into these membranes triggers their aggregation^47^. Our analysis also revealed that the largest HTTEx1Q72-mCherry puncta (volume > 5 µm^3^) were nearly all (90-100%) localized at the myotube surface at DCC 7-21 (Fig. 5c, e).

**Figure 5:**
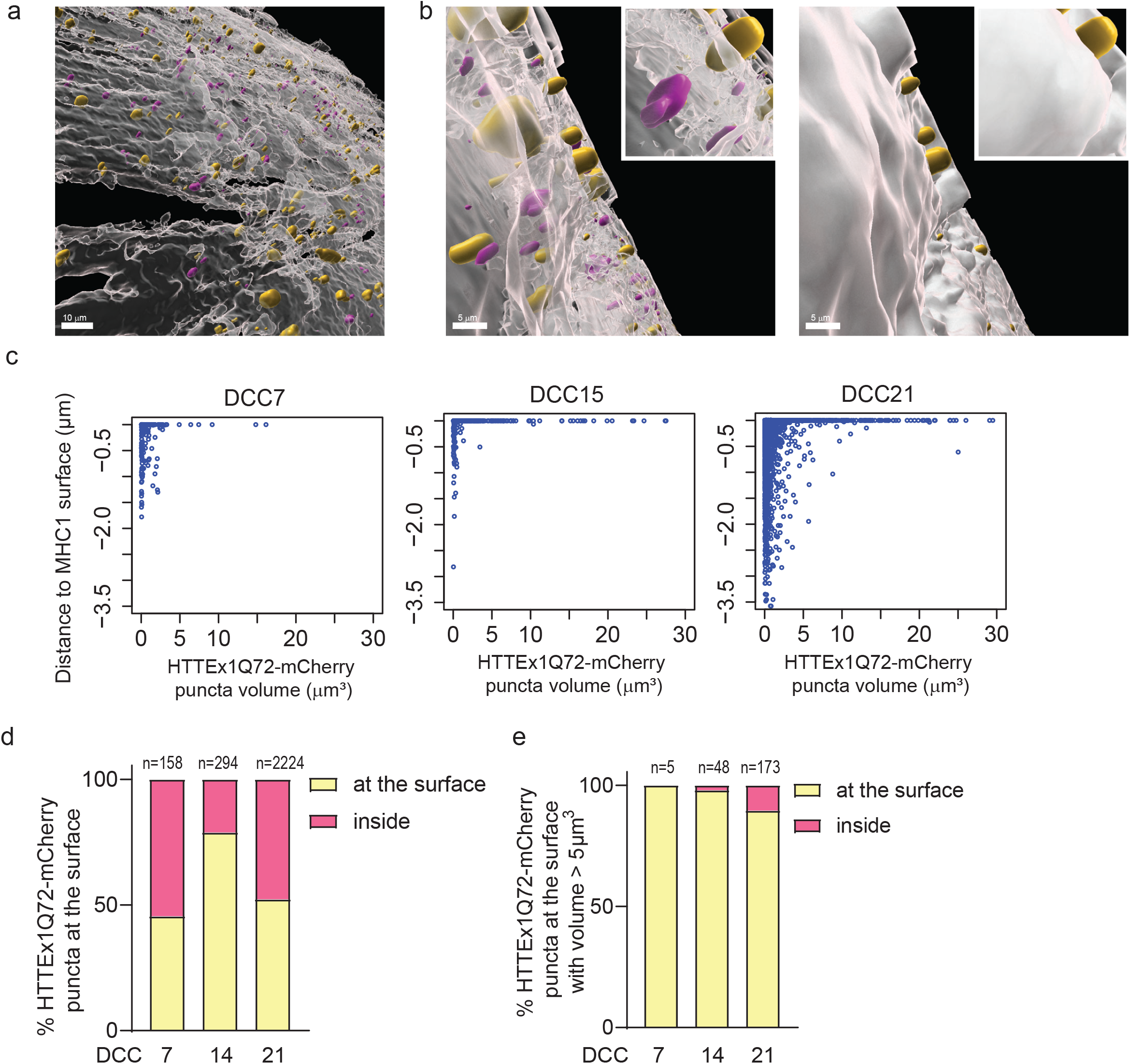
Overrepresentation of large HTTEx1Q72-mCherry assemblies at the MHC1+ myotube surface. **a**) Overview image of an Imaris model of MHC1+ Myo surface (white) with HTTEx1Q72-mCherry puncta associated with (yellow) and inside (magenta) the MHC1+ surface in Neu HTTEx1Q72-mCherry cl#75/Myo co-cultures. **b, left panel**) Zoom-in image of **(a)** showing yellow puncta penetrating the MHC1+ surface and magenta puncta not in contact with the surface. **b, right panel**) Same image as in “left panel” with non-transparent MHC1+ surface visualizing only the yellow puncta on the outside of the MHC1+ surface. **c**) Dot plots of HTTEx1Q72-mCherry puncta distance to MHC1+ surface against their volume at DCC 7 – DCC 21. **d**) Percentage of HTTEx1Q72-mCherry puncta at the MHC1+ surface at DCC 7 – 21. **e**) Percentage of HTTEx1Q72-mCherry puncta with a volume larger than 5 μm^3^ at the MHC1+ surface at DCC 7 – 21. The numbers (n) indicate the total number of puncta analyzed.

### HTTEx1Q72-mCherry transmission induces and aggravates pathological alterations in myotubes

An important open question in the field of misfolded proteins is whether the transmission can trigger or aggravate the pathology caused by the cell-autonomous presence of the toxic protein. Addressing this critical question will reveal whether toxic protein transmission is a novel disease pathway in neurodegenerative PMDs. Therefore, we assessed HD-specific pathological alterations in myotubes in the following Neu Myo co-culture combinations: 1) Neu control (ctr)/Myo ctr (no expression of the pathogenic HTTEx1Q72), 2) Neu ctr/Myo HTTEx1Q72-mCherry (cell-autonomous), 3) Neu HTTEx1Q72-mCherry/Myo ctr (transmission) and 4) Neu HTTEx1Q72-mCherry/Myo HTTEx1Q72-mCherry (transmission + cell autonomous).

Mitochondrial dysfunction is a characteristic observed in skeletal muscle obtained from HD patients and animal models^24^. Typically, a disbalance in the fission and fusion events occur which lead to more fragmented structures and a reduced filamentous network^34,48^. To assess the effect of cell-autonomous and transmitted HTTEx1Q72 on mitochondria fission/fusion we compared mitochondrial length, area weighted form factor and form factor in cell-autonomous and transmission co-cultures and compared these to control. We used MFDs to avoid contamination with neuronal mitochondria. We did not analyze cell autonomous + transmission co-cultures, where we cannot discriminate myotubes which received HTTEx1Q72-mCherry from neurons from those which did not. We observed a significant reduction in all three parameters when HTTEx1Q72-mCherry was expressed in myotubes and when the myotubes received HTTEx1Q72-mCherry from the neurons (Fig. 6a). A fragmentation of the mitochondrial filamentous network is likely to impair mitochondrial function. For example, in HD patient skeletal muscles, a reduction in ATP production has been observed and patients suffer from exercise-induced muscle fatigue already at preclinical stages of the disease^33,35^. To assess whether any functional change occurred in myotubes, we measured the myotube contractions. Strikingly, we observed a nearly complete loss of myotube contractions both measured by activity and contraction area, selectively in <u>transmission</u> or in <u>transmission + cell autonomous</u> co-cultures, despite the fact that these neurons displayed AP firing upon current injections (Fig. 6b, 2d). Interestingly, we also observed a smaller in magnitude but significant increase in activity in myotubes with cell autonomous expression that was independent of neuronal transmission. (Fig. 6b).

**Figure 6:**
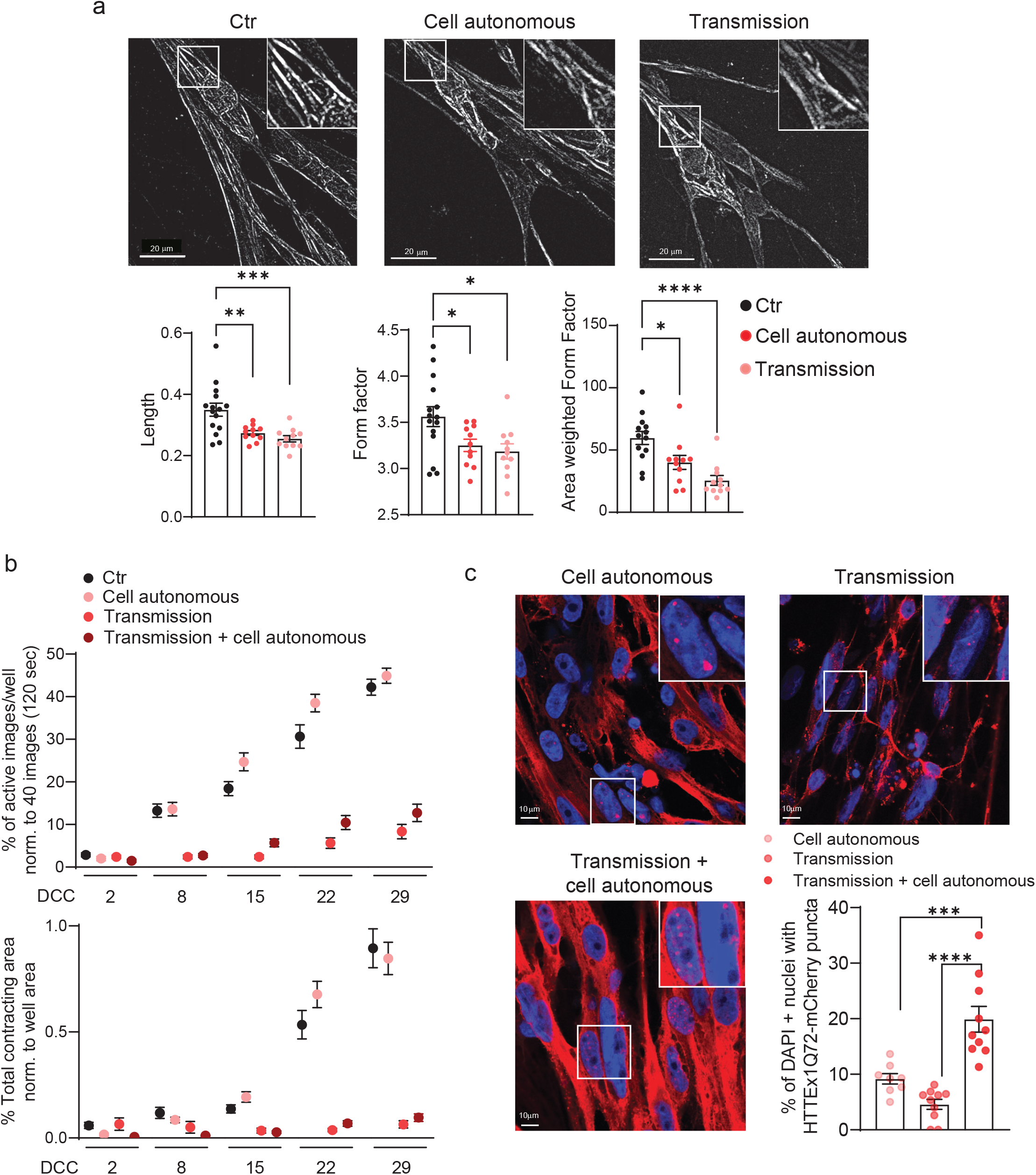
HTTEx1Q72 neuromuscular transmission causes structural and functional pathological alterations in myotubes in a dose-dependent manner. **a)** Images showing mitochondria marker TOMM20 mask outlining the mitochondria in myotubes in: <u>control</u> (Neu Ctr / Myo Ctr), Myo <u>cell-autonomous</u> (Neu Ctr / Myo HTTEx1Q72-mCherry) and <u>transmission</u> (Neu HTTEx1Q72-mCherry / Myo Ctr) co-cultures. Associated graphs show the quantification of structural parameters of the mitochondria in these mixed-genotype co-cultures. (n=30-40 images/genotype from 3 independent MFD co-cultures) **b)** Quantification of myotube contraction parameters measured in: <u>control</u>, <u>transmission</u>, <u>cell autonomous</u> and <u>transmission + cell autonomous (Neu</u> HTTEx1Q72-mCherry<u>/ Myo</u> HTTEx1Q72-mCherry) co-cultures at DCC 2 – 29 (n=22-27 wells/genotype and time point from 3 independent co-cultures). Three-way mixed design ANOVA: Significant transmission-dependent decrease in myotube contraction parameters dependent on time and independed of cell autonomous expression (p= 5.71e-34 upper panel, p=2.00e-21 lower panel). Significant increase in % of active images in cell autonomous cultures, dependent on time and independent of transmission (p-value=0.003) **c)** IF images of: cell <u>autonomous</u>, <u>transmission</u> and <u>transmission + cell autonomous</u> co-cultures at DCC 21, showing HTTEx1Q72-mCherry labeling in myotube nuclei (labelled with DAPI (blue)). Bottom right panel: percentage of myotube nuclei with HTTEx1Q72-mCherry puncta in the different genotype co-cultures. (n=10 images/genotype from 3 independent co-cultures) * = p ≤5 0.05; ** = p ≤5 0.01; *** = p ≤5 0.004; **** = p ≤5 0.0001 (one-way ANOVA, Dunnett’s correction). Abbr: **MFD** = microfluidic device. All averaged data are shown as the mean ± s.e.m.

To further assess pathological consequences of transmitted HTTEx1Q72 for myotubes we analyzed the extend of nuclear accumulation of HTTEx1Q72-mCherry aggregates in mixed genotype co-cultures. Nuclear aggregates in skeletal muscle of R6/2 mouse models of HD have been detected and their increase correlated with the worsening of disease pathology^36^. We observed the <u>lowest</u> number of nuclear aggregates in <u>transmission</u> co-cultures, this number was <u>slightly higher</u> when the protein was expressed <u>cell autonomously</u> in myotubes and <u>significantly increased</u> in the concurrent presence of <u>cell autonomous expression and transmission</u> from neurons (Fig. 6c).

### Mutant HTTEx1 is transmitted from the motor cortex to skeletal muscle *in vivo* in mice

With the *in vitro* experiments we so far demonstrated that mutant HTTEx1 is transmitted across the NMJ from neurons to muscle cells and that this induces pathological changes in the receiving myotubes. To understand whether this is also likely to occur in patients we studied the transmission of mutant HTTEx1 from the motor cortex to the skeletal muscles *in vivo* in mice. To this end we designed adeno-associated viruses carrying a floxed HTTEx1Q138-v5 plasmid (AAV_LoxP-Q138-v5). We chose a longer (138) CAG repeat, since mice are more resistant to CAG repeat expansion than humans. We used a 9 amino acid long v5 reporter tag to further exclude that the longer mCherry and Cre tags that we used in *in-vitro* were the driving force of the mutant HTTEx1 transmission. The AAV_LoxP-Q138-v5 was injected in the right hemisphere of the primary motor cortex (PM1) of mice expressing Cre selectively in the layer 5 pyramidal neurons (nex-cre mice). We observed HTTEx1Q138-v5 expression in the motor cortex (Fig. 7a). After 6 Months we analyzed the brachial spinal cord and the Triceps forelimb muscle and found HTTEx1Q138-v5 positive aggregates (Fig. 7b, c). We observed higher number of aggregates in contralateral side (left), confirming transmission following neuronal connectivity (Fig. 7d).

**Figure 7:**
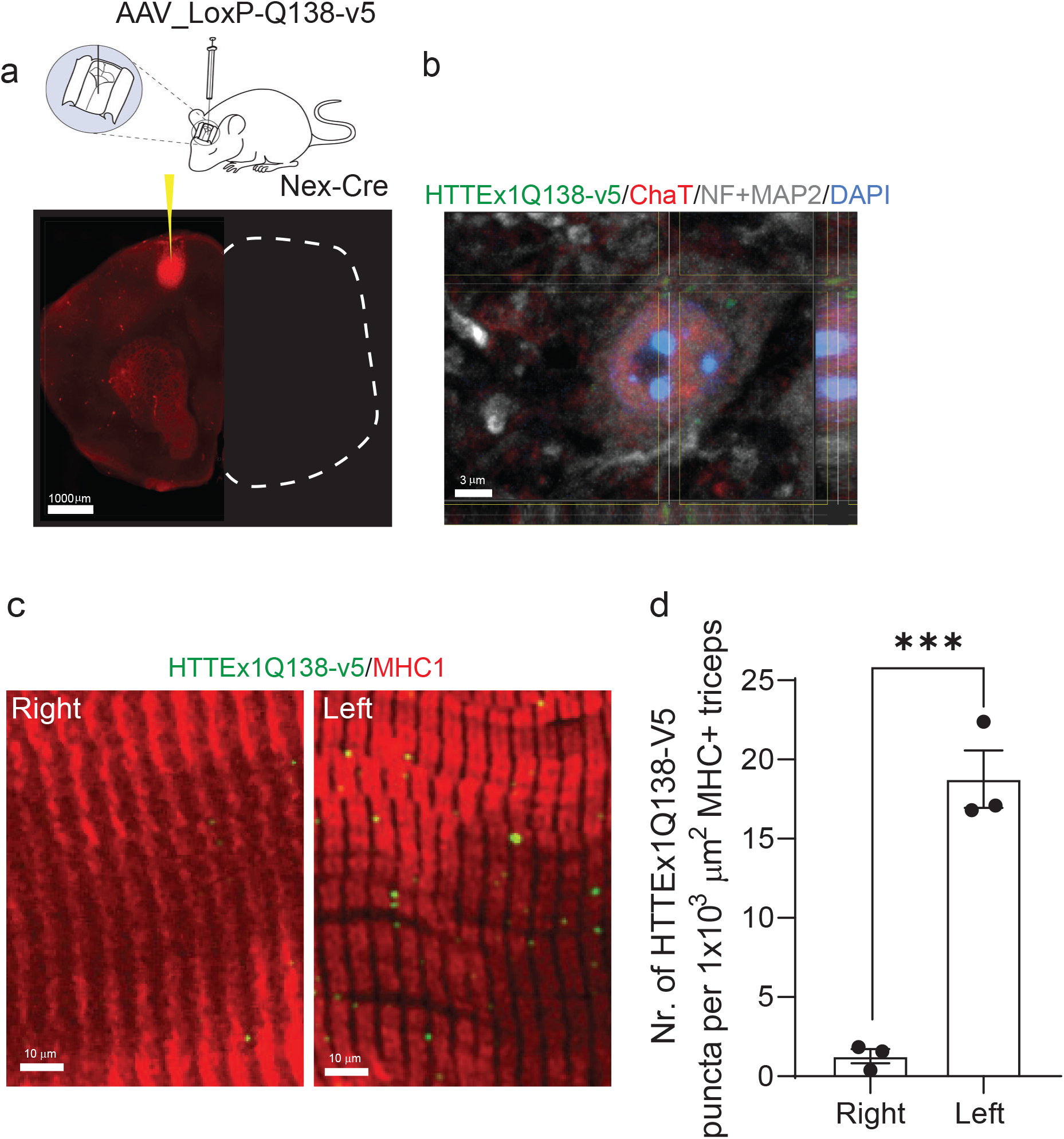
HTTEx1Q72 is transmitted from the motor cortex M1 to spinal motor neurons and skeletal muscles. **a, upper panel**) Image showing stereotactic injection of AAV_LoxP-Q138-v5 in mice expressing Cre specifically in projection neurons, including cortical pyramidal neurons. **a, lower panel**) IF image of HTTEx1Q138-v5 staining at the injection side in the M1 motor cortex. **b**) HTTEx1Q138-v5 puncta (detected with anti-V5 antibody) in ChaT+ motor neurons in the brachial spinal cord. **c**) IF of HTTEx1Q138-v5 puncta in the right and left triceps forelimb skeletal muscles stained with MHC1. **d**) Number of HTTEx1Q138-v5 puncta inside myotubes in right and left triceps (n=3, one data point corresponds to one animal and is a mean of 3-7 images, ***p=0.0007, Student’s t-test). Abbr: **ChaT** = choline acetyltransferase; **IF** = immunofluorescence; **MHC** = myosin heavy chain; **MAP2** = Microtubule-associated Protein 2; **NF** = neurofilament. All averaged data are shown as the mean ± s.e.m.

## Discussion

When cell-to-cell transmission of mHTTEx1 occurs and whether it is regulated by functional synaptic connectivity and can contribute to disease in an environment of ubiquitous expression of the mutant protein is to date not well understood. To advance our current understanding of these processes we established two *in-vitro* hiPSC derived neuro-muscular co-culture systems to study the role of neuromuscular connections in the development of HD-related skeletal muscle pathology. We provide evidence that neuromuscular transmission of mHTTEx1 can occur across the human neuromuscular synapse, likely already at early preclinical stages of disease, and contributes to skeletal muscle pathology. Furthermore, our findings suggest that mHTTEx1 transmission is more efficient when synaptic activity and density are increased. Finally, we show that mHTTEx1 is transmitted along the corticospinal pathway to skeletal muscles in mice *in vivo*. With the newly established cre-lox co-culture system we could follow HTTEx1Q72-cre transmission in the same culture, over weeks with fluorescent live-cell imaging. This revealed that transmission happened over time in the absence of detectable Em48+ aggregates in the neurons and that aggregation occurred first in the myotubes. Thus, mHTTEx1 is transmitted in the form of smaller protein structures, potentially as oligomers. Oligomeric structures are more soluble then larger aggregates, such as fibrils or inclusion bodies and therefore can diffuse more easily^49^. Furthermore, mHTTEx1 transmission happened already during the time of NMJ assembly *in vitro*. This thus may represent a very early pathological process in HD, possibly regulated by the immediate activity of the synaptic vesicle fusion machinery upon the contact of the growth cone with the muscle^50^.

Further, using Neu HTTEx1Q72-mCherry/Myo co-cultures we confirmed that transmission resulted in an increase of predominantly small protein-assemblies over time in the myotubes. HTTEx1Q72-mCherry puncta were associated with NMJ-forming AChR clusters types and transmission was positively correlated with NMJ density. Previously, it has been shown that synaptic density elevates neuron-to-neuron transmission of Tau^15^. These data together suggest that the density of synaptic connectivity between cells might be an important factor affecting toxic protein levels in postsynaptic cells. Additionally, our work encloses that mHTTEx1 secretion can be both regulated and constitutive. We found that HTTEx1Q72-mCherry transmission is elevated by neuronal depolarization, while preventing neuronal presynaptic release results in decreased transmission. This, together with previously published data on Aβ, Tau and mHTTEx1, strongly suggest that transmission occurs across functional synapses and is regulated by at least presynaptic activity^14-17,51^. In addition, a constitutive secretion is supported by our observation that hiPSC clones with higher HTTEx1Q72 expression levels showed more transmission to muscle (∼15-fold increase in HTTEx1Q72-mCherry puncta between myotubes cultured with clones #72 and #75). A positive correlation between mHTTEx1 concentration and aggregation has been previously shown^52^. Similar, for α-syn and Aβ a positive correlation between the intracellular levels and the amount that is released has been reported^53,54^. Thus, intracellular presence of misfolded proteins might trigger a highly sensitive stress response resulting in active transmission of the toxic species.

The pathobiological relevance of the misfolded protein transmission in an environment of ubiquitous expression of the toxic protein, as it is the case in HD, has not been assessed so far. It has been shown that transmission alone induces non-cell autonomous pathology in hiPSC-derived neurons, and *in vivo* in drosophila and C. elegans^14,17^. Furthermore, local presence of mHTTEx1 *in vivo* in mice and non-human primates’ results in propagation of the transgenic protein, and can induce motor deficits and cognitive decline after weeks/months (mice) or years (non-human primates)^12,55^. Here we demonstrate a causal link between transmission and pathology. Using the mixed-genotype co-culture system we dissected the contribution of transmission to the cell autonomous pathology. Strikingly, transmission of mHTTEx1 from neurons to myotubes, induced a severe decrease in myotube contractibility, which was not observed when the protein was expressed exclusively in myotubes. Huntingtin is a presynaptic protein and plays an important role in synaptic neurotransmission. Synaptic dysfunction is an early pathological phenomenon of HD^56^. Regulated release of mHTTEx1 might enrich its localization at the presynaptic site. Whether these factors contribute to the transmission selective loss of myotube contractions needs further investigation.

A transmission-selective pathology could be explained by a local increase of the toxic protein resulting from the specific route of transmission. With co-culture time we detected an increase of largest HTTEx1Q72-mCherry puncta, which preferentially localized to the myotube surface, suggesting that aggregation occurs at the myotube membrane. Supporting this observation, mHTT has a strong affinity for lipid membranes and bioengineered lipid bilayers have been shown to function as mHTTEx1 aggregate-promoting structures^57-61^. Aggregation of mHTT in the membrane is likely to cause a disruption of the lipid bilayer, with potentially a distorted localization of membrane receptors, including those required for normal transsynaptic signaling^62^.

Finally, we show that chronic inhibition of neurotransmitter release by exposing NM co-cultures to TeNT not only reduced release, but also resulted in increased nuclear aggregate pathology in HTTEx1Q72-mCherry expressing neurons. Clearance of misfolded proteins by the ubiquitin-proteasome system and autophagy is crucial to prevent protein accumulation to avoid aggregation^63,64^. Our finding suggest that toxic protein release might resemble a so far undefined pathway of misfolded protein clearance. A similar observation has been made for Aβ^53^. In this light, the new drug discovery strategies should promote release and prevent uptake of mHTT. The current antibody-based therapy designed to prevent Aβ, tau and α-syn accumulation in tau-and synucleinopathies, would also be a valuable strategy to test in HD^65,66^.

Taken together, the positive correlation that we observe between NMJ density and the HTTEx1-mCherry puncta, with transmission-triggered pathology suggests that the high number of synaptic connections in the CNS and between the spinal motor neurons and skeletal muscle makes these structures particular vulnerable to HD^35^. Given the peripheral phenotype, these findings also provide novel opportunities for biomarker development to assess the presence and contribution of this pathway in HD patients.

In a broader context, transsynaptic transmission of misfolded proteins is likely a common mechanism in PMDs, by which these toxic species spread through the brain and the periphery, contributing to a temporal decline of patient’s functional abilities.

## Materials and Methods

### iPSC culture and characterization

hiPSCs were previously generated from healthy adult human dermal fibroblast lines from a 32-year-old female from Invitrogen (C-013-5C), as described before (31). In brief, hiPSCs were maintained on Matrigel (354277, Corning) coated dishes with mTeSR 1 medium (05851, Stemcells Technologies) supplemented with Pen/Strep 1% (15070-063, ThermoFisher). Before differentiation, hiPSCs were confirmed to be pluripotent by western blot with OCT4 pluripotency marker (Supplementary Fig. 1c).

### Generation and differentiation iND3 Neurons

Neuronal differentiation protocol is described in Russell et al. (31) with smaller modifications. Briefly, hiPS cells were plated on matrigel in proliferation medium composed of DMEM/F12 with Glutamax (10565-018, Gibco) supplemented with 2% B27(17504-044,ThermoFisher) and 1% N2 (17502-048,ThermoFisher), 1% Pen/Strep (15070-063, ThermoFisher) supplemented with 10 ng/ml hEGF (PHG0315, ThermoFisher), 10 ng/ml hFGF (CTP0263, Invitrogen), with 10 µM Rock inhibitor (RI) for 1 day and 1 µg/ml doxycycline for 3 days, then progenitors were kept frozen in Cryostor freezing medium (07930, STEMCELL technology) or replated for immediate experiments.

### Generation of *MyoD* hiPSCs and differentiation to iMD3 myoblasts

Human MyoD cDNA was synthesized using sequence information from the Ensembl database (Accession number NM_002478) and cloned under the control of TRE tight (Tetracycline Response Element) promoter in a PiggyBac/Tet-ON all–in-one vector^67^. This vector contains a CAG rtTA16 cassette allowing constitutive expression of Tet-ON system and an Hsv-tkNeo cassette for generation of stable IPS clones. Genereation of MyoD hiPS was performed following a previously published protocol^37^. Briefly, 1 × 10^6^ hiPS cells were nucleofected by Amaxa nuclefector device using Human Stem Cell Nucleofector® Kit 1 (VPH-5012, Lonza) and program B-016 with 4 μg of MyoD plasmid and 1 μg of the dual helper plasmid. Subsequently cells were replated on matrigel plates with NutriStem medium containing 10 μM of Rock inhibitor. Antibiotic selection (G418 0.1 mg/ml) was applied after 48 hours. Stable clones appear within 1 week.

*MyoD* hiPS cells are seeded on 5µg/ml laminin-521-coated (Biolamina) in 5% KSR medium composed of Alpha-MEM (12571-063, Gibco), 5% KSR (10828028, Gibco), 1% Pen/Strep (15140-122, Gibco), 100µM β-Mercaptoethanol (21985-023, Gibco) + 1µg/ml DOX + 10 µM RI for 1 day. Medium change with 5% KSR medium + 1µg/ml Dox was done 24h later. 3 days after seeding, cells were frozen in Cryostor freezing medium or replated. Here, they are named iMD3 (hiPS-derived myoblasts Day 3). (Supplementary Fig. 2a).

### Plasmids generation

The human Huntingtin Exon1 carrying pathological 72 glutamines is fused to the Cre recombinase sequence (HTT_Ex1Q72-cre) or mCherry (HTT_Ex1Q72-mCherry) under the CAG promoter in a PiggyBac (PB) plasmid. A second PB plasmid is designed to carry a lox-stop-lox_GFP sequence (under the same CAG promoter). HTTEx1Q72-cre, HTTEx1Q72-mCherry, lox-stop-lox_GFP contructs were obtained by gene synthesis and cloned into PB backbone by Life Technology Europe BV. The three PB plasmids were nucleofected in hiPS Ngn2 or hiPS MyoD as described next.

### Generation of Cre-, mCherry and FloxP-stable lines

A single cell suspension of hiPS is collected upon Tryple Express Enzyme (12604-013, Gibco) detachment (5’ at 37°C). 1 × 10^6^ cells were resuspended in 100 μl of the nucleofection hESC solution 1 (Human Stem Cell Nucleofector® Kit 1/ Lonza #VPH-5012) where 5 μg of plasmids were added previously: 4 μg PB construct 1 μg Dual helper (expressing transposase). Nucleofection was performed using program B-016 on the Amaxa nucleofector II. Cells were immediately seeded after transfection into 6cm matrigel-coated dishes containing mTESR1 medium supplemented with 10 μM RI. 1µg/ml Puromycin selection is started 48-72h later. Clones were picked after 10 days. The clone was seeded in a new matrigel coated-35mm dish to amplify the new stable lines.

The stable lines were tested by temporal transfection with either the Q72-Cre construct in LoxP-GFP cell line and vice versa. Fluorescence was monitored daily with EVOS microscope to check their functionality. In parallel, the presence of HTTEx1Q72-cre or HTTEx1Q72-mCherry was checked via Western blot using Mab5492 antibody.

### From iMD3 to neuromuscular on-top coculture for live imaging

iMD3 cells were thawed (or replated) on laminin521-coated plates with 5% KSR medium+20ng/ml hFGF (CTP0263, Invitrogen) + 10µM RI for 3days with a seeding density of 2.5 × 10^6^ cells per laminin-521-coated 10cm dish. Medium change was done 24h later without RI. After 3 days from seeding, the 10cm dish was confluent. Cells were detached with Tryple Express Enzime (12604-013, Gibco), counted and seeded in medium C composed of DMEM F12-Glutamax + 5% FBS (SH30070.02, HyClone)+ 0.35% BSA (A1595, Sigma)+ 1%Pen/strep+ ITS 1:500 (354351, BD)+ 2µM CHIR99021 (1046, Sigma)+ 1µM Dorsomorphin (04_0024, Stemgent) + 1mM Dibutyryl-cAMP (BS0062, Biotrend) + 1µg/ml DOX + 10µM RI, and 2 × 10^5^ cells per well were seeded on laminin-521-coated 96 well IBIDI μ-plate (89626, IBIDI). RI and DOX was removed after 1 day. Medium was changed every other day until day 7 after seeding.

iND3 are thawed and seeded on top of the myotubes culture. 1.8 × 10^5^ iND3 are plated in neuronal differentiation medium composed of Neurobasal Medium (21103049, Thermofisher) + B27 with Vit. A (17504-044, Invitrogen) + N2 supplements (17502-048, Invitrogen) + Pen/Strep/Glutamax 1% supplemented with BDNF, GDNF, hNT3 (all from R&D at 10 ng/ml). Starting from day 2 of co-culture, medium change was done every other day. Neuromuscular cocultures were imaged at day 4, 7, 14, 21, 28 with Operetta (Perkin Elmer) in live-cell imaging (37 °C, 5% CO_2_) with 10x (NA 0.4) objective. GFP positive cells were counted manually.

### Electrophysiology

Neuromuscular co-cultures were established on glass in 24-well plates with a density of 3 × 10^5^hiPSC-derived iNgn2 (Neu HTTEx1Q72-cre or Neu HTTEx1Q72-mCherry cl.#75) neurons and 1.5 × 10^5^ hiPSC-derived iMyoD LoxP-GFP myotubes. The whole-cell patch-clamp technique was used to record action potentials of neurons at day of co-culture 7, 14 and 21. Co-cultures were taken from the incubator and transferred to the recording chamber with artificial cerebral spinal fluid (ACSF) containing (in mM): 125 NaCl, 25 NaHCO_3_, 2.5 KCl, 1.25 NaH_2_PO_4_, 2 MgCl_2_, 2.5 CaCl_2_ and 11 glucose, pH 7.4, constantly bubbled with 95% O_2_ and 5% CO_2_; 315–320 mOsm. The cells were kept at 30 - 32 °C and allowed to adapt for 20 minutes prior to recordings. Neurons were visualized with a LNScope (Luigs & Neumann) equipped with an oblique illumination condenser, a 60x objective (LUMPplanFI, NA 0.9) and a reflected illuminator (Olympus). Patch electrodes (5–7 MΩ) were pulled from borosilicate glass tubing and filled with an intracellular solution containing (in mM): 125 K-gluconate, 20 KCl, 10 HEPES, 10 EGTA, 2 MgCl_2_, 2 Na_2_ATP, 1 Na_2_-phosphocreatine, 0.3 Na_3_GTP, pH 7.2 (with KOH); 312.3 mOsm. Current-induced action potentials were recorded (with a holding potential of −70 mV) using a Multiclamp 700B amplifier (Molecular Devices) and digitized at 10 kHz. Recordings were performed at 30 – 32 °C in oxygenated ACSF. Igor Pro software (version 6.3, Wavemetrics) was used for both data acquisition and off-line analysis.

### Cell culture inserts

Culture-Insert 2 Well from IBIDI (81176) were used to seed iND3 and iMD3 in spatially separated areas of the well. Cell densities were adapted to this format. iND3 were seeded on Poly-L-lysine (P1524, Sigma) and laminin-521, iMD3 were seeded on laminin-521. Cells were monitored with EVOS M7000 microscope at day 1, 4, 8, 15 after the removal of the culture insert to check for the presence of GFP cells.

### Contractility assay in on-top co-culture

The primary readout for the amount of contraction in any on-top co-culture was captured as the total amount of motion within any given field of view over time. The Yokogawa CV7000 microscope with a 10x objective (NA 0.3) was adopted for this assay. The raw images were acquired as a series of 2560×2180 16-bit grayscale brightfield images with a frequency of 2 Hz., for a total amount of 60 images per field with 4 fields of view in each well. This assay was performed in live cell-imaging conditions (37°C, 5%CO_2_). At least 3 wells per experimental condition were acquired and analyzed.

For each consecutive pair of image frames a motion field was computed which provides, for each pixel location, a direction and magnitude of projected spatial motion. Thus, for N image frames we obtained N-1 motion frames. A numerical threshold on the magnitude of the motion vectors was applied to eliminate possible noise and vibration artefacts and to obtain a reliable binary image map of region-of-contraction. The union of all such pixels over all motion frames in the time series was computed and used as the final region-of-contraction map for comparative analysis between cell lines or treatments. These values were used to describe the on-top co-culture functionality as follows:

- Total contracting area normalized to well area (%): sum of moving pixels normalized to the acquired fields area (namely, sum of the pixels occupied by the 4 fields of view)
- Active images per well normalized to total number of images (%) where active image are images for which a pixel movement was detected

### Antibodies and dyes

For western blot: 1:500 Embryonic myosin (MHC3) (F1.652, DHSB), 1:500 Postnatal myosin (MHC8) (N3.36, DSHB), 1:1000 Islet 1(AF1837-SP, R&D), 1:2000 Doublecortin (DC) (4604, Cell signaling), 1:1000 Choline acetyltransferase (ChAT) (AB144P, Merck) 1:250 Oct4 (09-0023, Stemgent), 1:5000 GAPDH (ab9485, abcam), 1:5000 β-actin (A5441, Sigma), 1:5000 MAB5492 (MAB5492, Sigma-Aldrich).

For immunofluorescence: Hoechst 33342, 5µg/ml α-bungarotoxin (B1196, Thermofisher), 1:2000 Bassoon (141 013, Synaptic Systems), 2µg/ml Tetanus neurotoxin (T3194-25UG, Sigma), 1:1000 Neurofilament M (171 204, Synaptic systems); 1:1000 Myosin Heavy Chain 1 (05-716, Millipore), 1:5000 mCherry (ab205402, Abcam) 1:500 TOMM20 (ab186735, Abcam), 1:500 EM48 (MAB5374, Merck), 1:100 ChAT (AB144P, Merck), 1:2000 Map2 (ab5392, Abcam), 1:800 V5 (D3H8Q, Cell Signaling), 1:40 NF2H3 (AB2314897, DSHB).All the secondary antibodies were Alexa conjugated from Jackson ImmunoResearch and used 1:1000 for 1 h at RT.

### Immunocytochemistry for GFP+ cells

Cells were fixed in 4% paraformaldehyde (PFA) at room temperature for 7 min, followed 3 DPBS (14190, Sigma-Aldrich) washings, five minutes each. DPBS supplemented with 0.1% Triton X-100 (for permeabilization) and 1% BSA (blocking) was used for primary antibodies labelling, overnight at 4°C. After three washing steps with DPBS, cells were incubated for 1h with secondary antibodies (Invitrogen). Afterwards, cells were washed in DPBS and incubated with Hoechst 33342 in ddH2O for 10min. Ibidi mounting medium (50001, IBIDI) is added to the wells and stored at 4°C. Images were acquired with LSM900 microscope with Plan-Apochromat 63x/1.40 Oil DIC M27, using Zen 3.2 (Blue edition) software.

### Western Blot

IPSC-derived neurons, myotubes or co-culture cells were harvested at different time points, washed twice with ice-cold PBS, and subsequently lysed in RIPA buffer supplemented with complete EDTA-free protease inhibitor mixture (11873580001, Roche). Lysates were incubated on ice for 15 min and cleared via centrifugation (10,000 × g) for 10 min at 4 °C. Supernatants were collected, and the protein concentration was determined using a BCA assay kit (Thermo Scientific Pierce, 23227). Lysates were resolved using standard SDS-PAGE gels and after blocking, blots were incubated with primary antibodies overnight at 4 °C. After washing, blots were incubated with secondary antibodies and visualized using SuperSignal Femto chemiluminescent detection kit (Thermo Scientific) in Odyssey Infrared Imager (LiCor, 9120). The image in Fig. 1b is representative from 3 independent experiments.

### Immunofluorescence on coverglass culture

Cells on glass coverslips (in format 24 well plate with 3 × 10^5^ myotubes and 3 × 10^5^ neurons density) were fixed for 5 min in 4% PFA/4% sucrose at RT, permeabilized with PBS+/+ (D8662, Sigma, supplemented with 1 mM MgCl2 and 0.1 mM CaCl2)/Triton-0.1%, blocked with 5% BSA in PBS+/+ and labeled with primary antibodies in PBS+/+ (D8662, Sigma) and 5% BSA overnight at 4°C and secondary antibodies for 1h RT. PBS+/+ washes were performed after each antibody incubation. Coverslips were mounted on glass slides in Prolong (P36930, Invitrogen).

### Microfluidic devices (MFD) culture

We used XonaChips XC450 devices from Xona Microfluidics. iMD3 cells were growth in 5%KSR medium (Alpha-MEM (12571-063, Gibco); 5% KSR (10828028, Gibco); 1% Pen/Strep (15140-122, Gibco); 100µM β-Mercaptoethanol (21985-023, Gibco), supplemented with 1µg/ml doxycycline (D1822, Sigma) and 20ng/ml FGF (300-112P, Gemini Bio). At DIV 3 the cells were seed in final format for differentiation: In myocytes side, 3 × 10^5^ cells were seed and in the neuronal side, 1.5 × 10^5^ cells in 5 µl medium, were seeded to give support to the motor neurons. The myotubes growth for 7 days in “Differentiation medium” (DMEM F12-Glutamax (10565-018, Gibco); 5% FBS (SH30070.02, HyClone); 1:500 ITS (354351, BD); 0.1% BSA (A1595, Sigma); 1% Pen/Strep, supplemented with 2µM CHIR99021 (1046, Sigma); 1 µM Dorsomorphin (04_0024, Stemgent); 1mM Dibutyryl-cAMP (BS0062, Biotrend). Then, 3 × 10^5^ neurons were seeded in the neuronal compartment and the culture growth in “neuronal medium” (Neurobasal TM Medium; B27 (17504-044, Invitrogen); N2 supplement (17502-048, Invitrogen) 1% Pen/Strep/Glutamax; and BDNF, GDNF, hNT3 (all from R&D).

For immunofluorescence experiments, the culture was fixed at different time points for 10 min in 4% PFA/4% sucrose at RT, then the immunofluorescence protocol was followed.

### Mitochondrial Morphology Quantification

Mitochondrial shape parameters were quantified using the open-source software package ImageJ. Measuring Mitochondrial Shape with ImageJ. In: Strack S., Usachev Y. (eds) Techniques to Investigate Mitochondrial Function in Neurons. Neuromethods, vol 123. Humana Press, New York, NY. https://doi.org/10.1007/978-1-4939-6890-9_2) Briefly, images were background-subtracted (rolling ball radius = 50 pixels) and uneven labeling of mitochondria was improved through local contrast enhancement using contrast-limited adaptive histogram equalization (“CLAHE”). To segment mitochondria, the “Tubeness” filter was applied. After setting an automated threshold, the “Analyze Particles” plugin was used to determine the area and perimeter of individual mitochondria and the “Skeletonize” function was used to measure mitochondrial length.

Three parameters were assessed:

- Mitochondrial length: the length reports the mitochondrial length or elongation in pixel, after the mitochondria are reduced to a single-pixel-wide shape (“Skeletonize” function on ImageJ).
- Form factor (FF): The FF value describes the particle’s shape complexity of the mitochondria, as the inverse of the circularity.
- Area-weighted form factor (AWFF): a variant of FF with a bias towards larger mitochondria or mitochondrial networks. AWFF provides more realistic results in cases where highly elongated mitochondria are overlapping

### Animal husbandry

Adult NEX-Cre were kindly provided by Dr. Sandra Goebbels (Max-Planck-Institute of Experimental Medicine, Goettingen, Germany). All mice were housed in temperature (22°C) and light-controlled environment on a 12-light dark cycle and had access to food and water ad libitum. All experimental procedures were carried out according to Basel University animal care and use guidelines. They were approved by the Veterinary Office of the Canton of Basel-Stadt, Switzerland.

### Delivery of viral vectors

Four-week old female NexCre mice were anaesthetized by the administration of 4% isoflurane, were maintained under isoflurane anesthesia (1-2%) and kept warm with a heating pad (53800, Stoeling). The head was fixed to a stereotaxic frame (Kopf Instruments) with ear bars and the skin was disinfected with 70% ethanol and polyvidone iodine. The skin was cut with surgical scissors to expose the skull, allowing the identifications of bregma and lambda. Using a borosilicate glass pipette and a pressure ejection system (Eppendorf) 250 nl of the self-complementary AAV-9/2-DIO-mHTTExon1Q138-V5 (VVF, Zürich) were injected in the layer V of the primary motor cortex, using the following coordinates AP (anterior-posterior): + 1.18, ML (medial-lateral): + 2.00, DV (dorsal-ventral): + 2.00, according to the Paxinos and Franklin mouse brain atlas (Paxinos and Franklin, 2019, eBook ISBN: 9780128161609). The mice were placed in a recovery cage to awaken before returning to their home cage. 6-month-old female mice were anesthetized by the administration of 4% isoflurane and were fast decapitated.

### Immunohistochemistry for spinal cord and muscle samples

The spinal cord, biceps and triceps were dissected on ice and embedded in low-melting agarose (<u>16520050</u>, ThermoFisher Scientific). Samples were sliced in 100-150 µm thick sections using a vibratome (VT1200, Leica). Then, fixed in 4% paraformaldehyde (PFA) at room temperature for 10 min, followed by 3 DPBS (14190, Sigma-Aldrich) washings, 10 minutes each. DPBS supplemented with 0.1% Triton X-100 (for permeabilization) and 1% BSA (blocking) was used for primary antibodies labelling, for at least 2 days at 4°C. After three washing steps with DPBS, cells were incubated for 3h with secondary antibodies (diluted 1:800, Invitrogen). Afterwards, cells were washed in DPBS for 3 times. Sections were then mounted on glass slides using ProLong Gold (P10144, ThermoFisher Scientific) and acquired with LSM800 with 40x objective (ZEISS, EC Plan-NEOFLUAR 40X/1,3 Oil)

### Image acquisition and analysis

Fluorescence signals in “on top” culture for iPSC-derived co-culture were imaged with Zeiss LSM-700 system with a Plan-Apochromat 40 × /NA 1.30 oil DIC, using Zen 2010 software. For bin analysis in MFD, section of 0-160, 160-320 and 320-480 µm were taken in the using Zeiss LSM-800II inverted system with a Plan-Apochromat 40 × /NA 1.30 oil DIC, using Zen blue 2.6 software. Whole-cell, 16-bit stacks images with 0.33-μm step size were acquired (15–30 planes). Immersion oil with 1.518 refractive index at room temperature was applied to the lens. Coverslips were mounted with ProLong Gold anti-fade reagent (P36930, Thermofisher) with a refractive index of 1.46. All images were acquired with identical microscope settings within individual experiments. Brightness and contrast were adjusted equally for all images, and cropped insets were generated in the same manner among all the experiments to facilitate visualization of representative cells. Saturation was avoided by using image acquisition software to monitor intensity values. For any image adjustment, identical settings were always applied to all cells, irrespective of genotype.

Cells that were clumped or overlapping were excluded from quantification. For quantification, values were averaged over multiple cells from at least three independent culture preparation. Quantification of number and volume HTTEx1Q72-mCherry puncta was done using Imaris Software (v.9.6.0; Oxford Instruments) function and measurement based on mCherry fluorescence staining. Aggregates with volume above 0.02 and below 30 µm^3^ and localized within the surface generated based on MHC1 staining were analyzed.

Intracellular localization of aggregates was analyzed using distances between surfaces generated based on mCherry staining for aggregates and MHC1 staining for muscle by Imaris software. Localization at the surface was defined as distance between 0 and 0.05 µm.

Quantification of nuclei containing HTTEx1Q72-mCherry were done using Image J software. Images were background subtracted and after setting an automated threshold a mask for DAPI positive nuclei (in MHC1 positive myotubes or MAP2 positive neurons) was applied and the “Analyze Particles” plugin was used to determine the number of puncta per nucleus.

Quantification of HTTEx1Q138-V5 puncta in triceps were done using Image J software. Images were background subtracted and after setting an automated threshold, the “Analyze Particles” plugin was used to determine the number of puncta in MHC1 positive staining. Integrated density of EM48/MAP2+NF staining was done using Image J software. Images were background subtracted and after setting an automated threshold, integrated density was measured in the full image, to consider EM48 staining in the soma and neurites of the neurons. The values were normalized to the neuronal market MAP2 and NF.

AChR clusters were analyzed using Imaris Software (v.9.6.0; Oxford Instruments) based on immunofluorescent images acquired by Confocal microscope. 3D reconstruction of AChR and BSN structures were done using Imaris surface function. Automatically generated values for volume and sphericity were used to characterize the clusters. Only structures with volume above 0.02 and below 20 µm^3^ were analyzed for BSN and only structures above 0.024 µm^3^ for AChR. Distances between the surfaces provided by Imaris software were used to identify AChR cluster and BSN association. Association was defined as distance below 0.05 µm.

Multiple images were analyzed using Imaris Batch function. The data on volume, sphericity and distances between the surfaces were exported and further analyzed using R (v.4.0.5; https://www.R-project.org/) and RStudio software (v. 1.4.1106, https://www.rstudio.com/), using base R and ggplot2 (v.3.3.5).

## Statistical analysis

Data analysis was performed with GraphPad Prism version 8.0 (GraphPad Software, La Jolla, CA) and using R software (v.4.0.5; https://www.R-project.org/). Individual data sets were tested for normality with the Shapiro-Wilk, D’Agostino & Pearson or Kolmogorov-Smirnov test. Statistical significance of differences between groups was assessed by unpaired or paired two-tailed Student’s t-test or ANOVA as indicated. Time series experiments were analyzed using Mixed Design ANOVA or Mixed Effect Linear Model when missing data were present. For data with non-normal distribution the non-parametric Wilcoxon rank sum or Kruskal-Wallis tests were used. For comparison on group proportions Chi square test or Fisher’s exact test (for samples with expected frequencies below 5) were used. p-values < 0.05 were considered significant. For analysis of variables relationship simple linear regression and Pearson correlation were used. Data are presented as mean ± standard error of the mean (s.e.m.). All statistical tests and results are reported in Supplemental Table 1.

## Data availability

Data that support the findings of this study are available from the corresponding author upon reasonable request.

## Code availability

Custom code used in this study is available from the corresponding author upon reasonable request.

## Acknowledgements

We thank Patricia Valerio from the Department of Biomedicine (DBM), University of Basel, for providing us training with the stereotactic injection technology. We also thank the DBM and Biocenter imaging and bioinformatic facilities, University of Basel for their expert input. Furthermore, we thank Inga Galuba, Isabelle Claerr and Dr. Gianluca Santarossa, Novartis Institute of Biomedical research (NIBR), Basel for their support with the Yokagawa imaging system. We thank Prof. Josef Bischofberger, DBM, university of Basel, for his support and discussions. This work was supported by a Swiss National Science Foundation professorship grant (PP00P3_163937; PP00P3_194806) and a Synapsis foundation – Alzheimer Research Switzerland ARS grant for principal investigators to EPV.

## Author information

These authors contributed equally: Margarita C. Dinamarca and Laura Colombo

## Novartis Institute for Biomedical Research, Basel, Switzerland

Isabelle Fruh, Imtiaz Hossain, Daniela Gabriel & Matthias Müller

## Contributions

M.C.D. and E.P.V. conceptualized the study. M.C.D., L.C., N.E.T., A.E., M.M. and E.P.V. developed the methodology. I.H and D.G. developed the mathematical algorithm. M.C.D., L.C., N.E.T., A.G., I.F. and E.P.V. performed the experimental investigations. M.C.D., L.C., U.B., A.G. and E.P.V. performed the data analysis. M.C.D. and U.B visualized and curated the data. E.P.V. wrote the original draft. M.C.D., U.B. and E.P.V. wrote and edited the manuscript. All authors edited and/or reviewed the manuscript. E.P.V. supervised the project and acquired funding.

## Ethics declarations

Competing interests

The authors declare no corresponding interests.

