## Supplemental Figures and legends for "Transmission-selective muscle pathology induced by active propagation of mutant huntingtin across the human neuromuscular synapse"

Supplemental Figure 1

**a, left panel**) Schematic of a plasmid map used for generating the three proneuronal HTTEx1 expressing iNgn2 hiPSC lines. **a, right panel**) Schematic of a plasmid map used for generating the two promuscle iMyoD hiPSC lines. The iMyoD line is isogenic to the iNgn2 line. **b**) WB of HTTEx1Q72 expression visualized with Mab5492 Ab in hiPSCs HTTEx1Q72-mCherry clones: 71, 72 and 75. To visualize the expression of HTTEx1Q72 in the hiPSC HTTEx1-Q72-cre line exposure time of the western blot had to be increased (right). (n=3) **c**) Representative WB of pluripotent marker Oct4 in iNgn2; HTTEx1Q72-cre hiPSCs and precursor neurons at DIV 1. **d**) Quantification of Em48 fluorescent intensity in neurons at DIV7 and 21, **$p<0.01$, ***$p<0.0001$ (5 pictures from 3 independent cultures, one-way ANOVA, Tukey's correction). **e**) IF images visualizing absence of GFP expression in iMyo;LoxP-GFP hiPSCs not transfected with HTTExQ72-cre plasmid (left image) and presence of GFP expression for the cells transfected with HTTExQ72-cre plasmid (middle image) and expression of mCherry when the hiPSC iNgn2;HTTEx1-cre are transfected with a LoxP-mcherry plasmid (Right image). Abbr: **DIV** = day in vitro; **IF** = immunofluorescence; **Int. density** = integrated density; **WB**= western blot. All averaged data are shown as the mean ± s.e.m.

Supplemental Figure 2

**a**) Diagram showing the 2-step protocol used to establish the neuromuscular co-culture. Step 1: iMyoD and iNgn2 hiPSCs are prior to assembly separately cultured and differentiated by exposing them for 3 days to DOX, into myoblasts and precursor neurons, resp. Myoblasts are prior to co-culture seeded and cultured in myotube promoting media to allow further differentiation and maturation of the myoblasts into myotubes. After 10 days the precursor neurons are seeded on top of the myotubes. This we indicate as DCC0. The co-culture is grown in motor neuron promoting media. **b**) Voltage traces showing examples of the different current-induced AP firing patterns recorded in Neu HTTEx1Q72-cre neurons when voltage clamped at -70 mV. **c, left panel**) Images showing area of contraction in HTTEx1Q72-cre / Myo LoxP-GFP before and after treatment with 2.5 μM BgTx at DCC22. **c, right panel**) Quantification of the % total contracting area normalized to well area at DCC 22, before and after treatment with BgTx *** = $p=0.0005$ (Wilcoxon matched-pairs signed rank test). **d**) Quantification of the variability in the number of active images (= images with myotube contractions) (upper graph) and the variability in contracting area (lower graph) between the four different fields recorded in one well at progressive days of co-culture (n=3-9 wells/time point in 1-3 independent cultures, Two-way Mixed ANOVA (without DCC2 due to missing values): time dependent significant difference in variability in activity (upper panel) between co-culture and muscle ($p=9.09e-8$) and time independent significant difference in total contracting area (lower panel) between co-culture and muscle ($p=1.77e-07$). Post-hoc one-way repeated measures ANOVA: significant decrease with time in variability in activity (left panel) for co-culture (p=0.0004), and no significant effect for muscle. Abbr: **ms** = milliseconds; **BgTx** = α-bungarotoxin (labels AChRs); **DCC**: day of co-culture. All averaged data are shown as the mean ± s.e.m.

Supplemental Figure 3

**a)** Time-line of co-culture experiment where Neu HTTEx1Q72-cre and Myo LoxP-GFP are seeded in separate inserts which are placed in the same well to avoid physical contact between the two cell lines, but ensure exposure of the two cell lines to the same culture medium. **b, top schematic)** Depicting 1 well with the inserts. The insert with neurons is coated with PLL and Lam, the one with myotubes are seeded with Lam. The space between the inserts is not coated to avoid movement and attachment of the cells to this region. **b, bottom, left panels**) Bright field image showing physical separation of Neu HTTEx1Q72-Cre and Myo LoxP-GFP cells (left). Right panel shows absence of GFP+ myotubes in these cultures at DCC4 - 21 (n=4 independent devices). **b, bottom, right panel**) IF images to show expression of the neuron-specific marker NF200, myotube-specific marker MHC1 and the nuclear marker Hoechst. Abbr: **PLL** = Poly-L-Lysine; **Lam** = Laminin.

Supplemental Figure 4

**a, left image**) IF labeling of neuromuscular co-culture in MFD to visualize the NMJs (appositions between α-BgTx and BSN) between Neu HTTEx1Q72-mCherry cl#75 and Myo LoxP-GFP. The Image is taken from the myotube compartment. **a, middle image**) Same image as in left, with transparent surfaces for MHC1, AChRs and BSN. The surface was created with the 'surface' function in Imaris. **a, right image)** Zoom-in of middle image to visualize the AChR clusters, and an apposition of an AChR cluster with BSN (yellow arrowhead). **b)** Volume against sphericity of the AChR clusters based on Imaris surface measurements. The red dotted lines indicate thresholds used to define four cluster classes - 20 $\mu m^3$ for the volume and 0.6 for the sphericity. Data points for clusters with close appositions between AChRs and BSN (<0.05 µm distance between the surfaces) are colored in blue. **c)** Pie chart of the proportions of four cluster classes based on data in panel (b) (Data from 12 images of Neu HTTEx1Q72-mCherry cl#72 and Myo LoxP-GFP co-cultures in MFD.). **d)** Distribution of four AChR cluster types in percentage, found on Myo when cultured with or without neurons. **e)** Comparison of the numbers of AChR clusters in four classes found on Myo when cultured with or without neurons. Number normalized to muscle volume (based of MHC1 surface). One data point corresponds to one image (12 images analyzed per condition). ** = $p \leq 0.01$; **** = $p < 0.0001$ (Student's t-test) Abbr: **AChRs** = acetylcholine receptors; **BgTx** = α-bungarotoxin (labels AChRs); **BSN** = Bassoon; **DCC** = days of co-culture; **IF** = immunofluorescence; **MHC** = myosin heavy chain, **NMJs** = neuromuscular junctions; **MFD**= microfluid devices. All averaged data are shown as the mean ± s.e.m.

Supplemental Figure 5

**a**) Quantification of the NFM area normalized to the total MHC1 area in each bin in MFD at DCC21. ** = $p \leq 0.01$; **** = $p < 0.0001$ (Kurskal-Wallis Test with Dunn's correction, n=15 pictures per bin from 1 experimental set) **b**) Number of HTTEx1Q72-mCherry puncta in myotubes counted at DCC 7 – 21, in bin 1, 2, and 3 in the myotube compartment of a MFD (n=3 independent co-cultures/time point, data point: mean of 5 images) **c**) Percentage of HTTEx1Q72-mCherry aggregates associated with AChR clusters at DCC 7 – 21. (n = total number of aggregates analyzed -written above each bar). Abbr: **AChRs** = acetylcholine receptors; **DCC** = days of co-culture; **MHC** = myosin heavy chain, **NFM** = neurofilament m; **MFD**= microfluid devices. All averaged data are shown as the mean ± s.e.m.

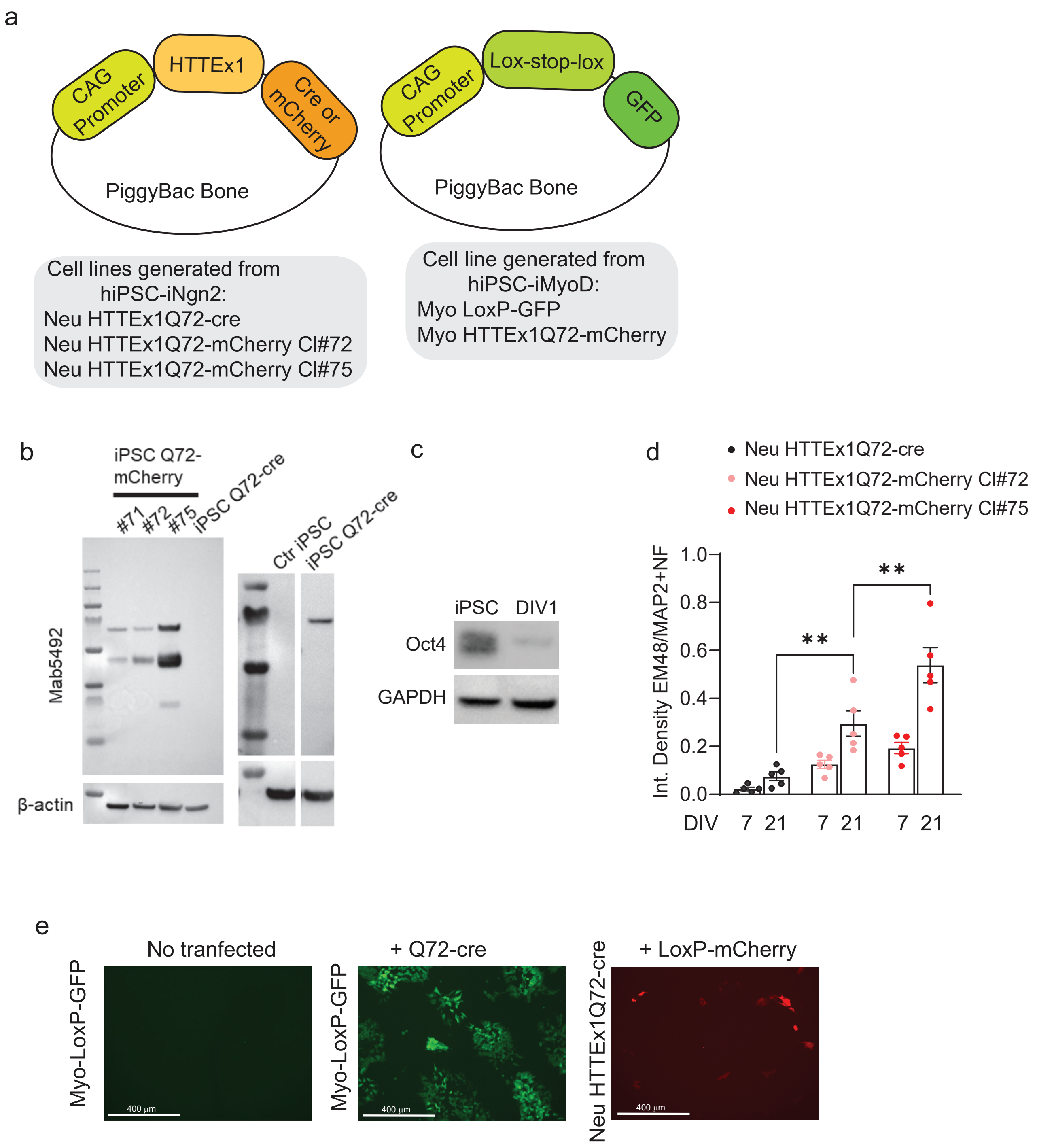

a
CAG Promoter
HTTEx1
Cre or mCherry
PiggyBac Bone
CAG Promoter
Lox-stop-lox
GFP
PiggyBac Bone
Cell lines generated from hiPSC-iNgn2:
Neu HTTEx1Q72-cre
Neu HTTEx1Q72-mCherry Cl#72
Neu HTTEx1Q72-mCherry Cl#75
Cell line generated from hiPSC-iMyoD:
Myo LoxP-GFP
Myo HTTEx1Q72-mCherry
b
iPSC Q72-mCherry
#71
#72
#75
iPSC Q72-cre
Ctr iPSC
iPSC Q72-cre
Mab5492
β-actin
c
iPSC
DIV1
Oct4
GAPDH
d
Neu HTTEx1Q72-cre
Neu HTTEx1Q72-mCherry Cl#72
Neu HTTEx1Q72-mCherry Cl#75
Int. Density EM48/MAP2+NF
1.0
0.8
0.6
0.4
0.2
0.0
**
**
DIV 7 21 7 21 7 21
e
No tranfected
+ Q72-cre
+ LoxP-mCherry
Myo-LoxP-GFP
Myo-LoxP-GFP
Neu HTTEx1Q72-cre
400 µm
400 µm
400 µm

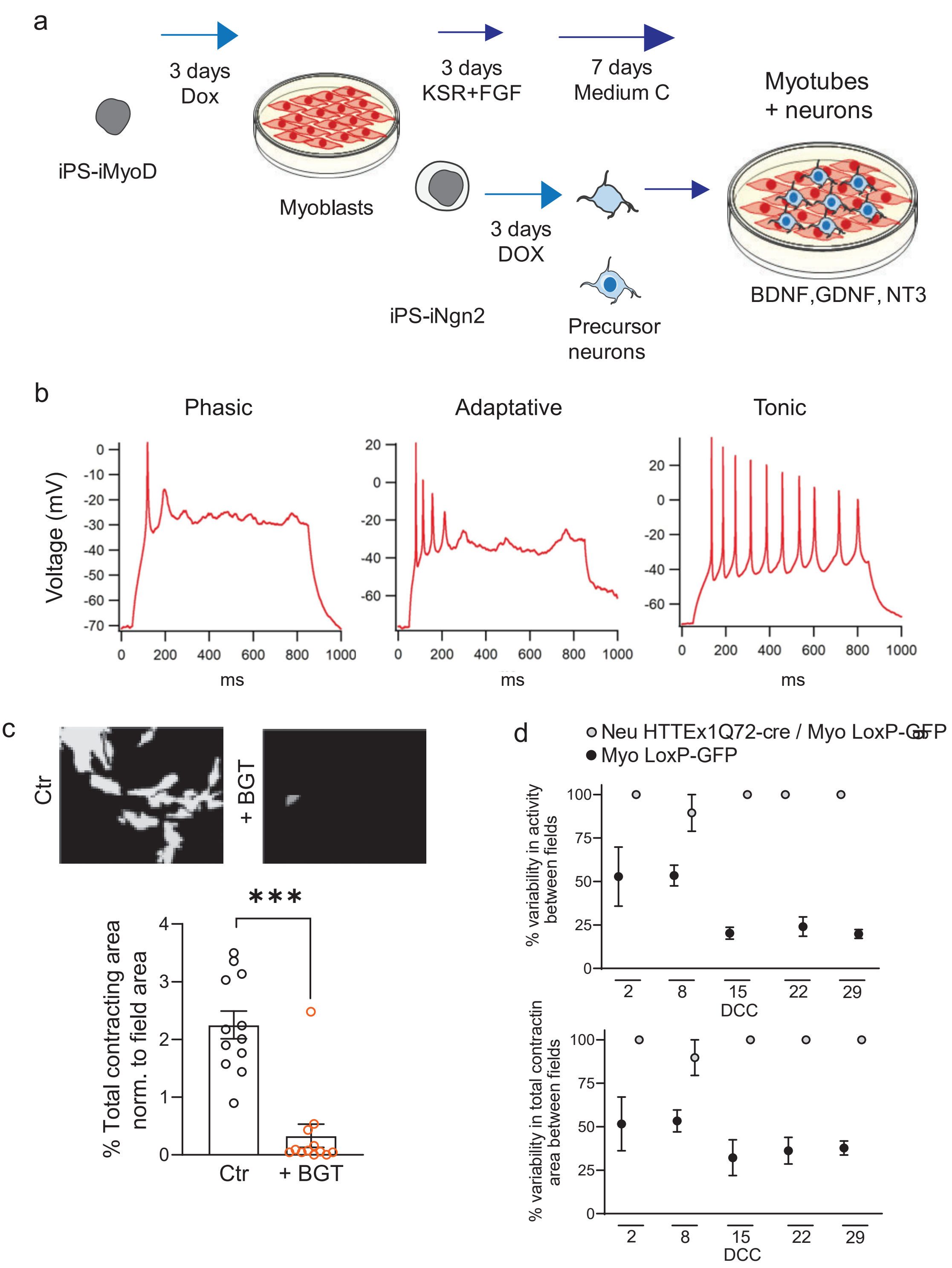
a
3 days
Dox
iPS-iMyoD
Myoblasts
3 days
KSR+FGF
7 days
Medium C
Myotubes
+ neurons
3 days
DOX
iPS-iNgn2
Precursor
neurons
BDNF,GDNF, NT3
b
Phasic
Adaptative
Tonic
Voltage (mV)
ms
c
Ctr
+ BGT
***
% Total contracting area
norm. to field area
Ctr
+ BGT
d
Neu HTTEx1Q72-cre / Myo LoxP-GFP
Myo LoxP-GFP
% variability in activity
between fields
DCC
% variability in total contractin
area between fields
DCC

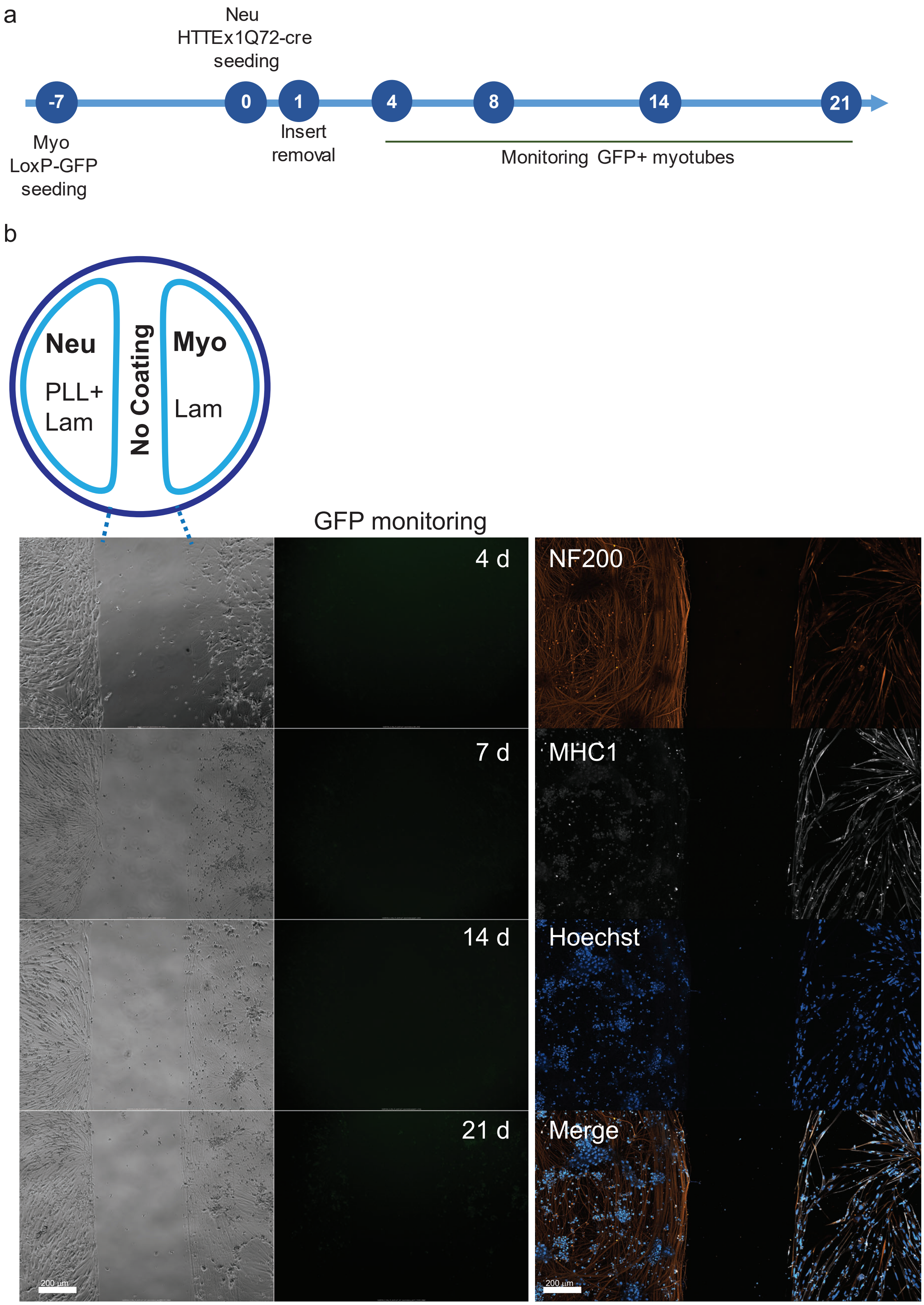
a
Neu
HTTEx1Q72-cre
seeding
-7
0
1
4
8
14
21
Myo
LoxP-GFP
seeding
Insert
removal
Monitoring GFP+ myotubes
b
Neu
PLL+
Lam
No Coating
Myo
Lam
GFP monitoring
4 d
7 d
14 d
21 d
NF200
MHC1
Hoechst
Merge
200 µm
200 µm

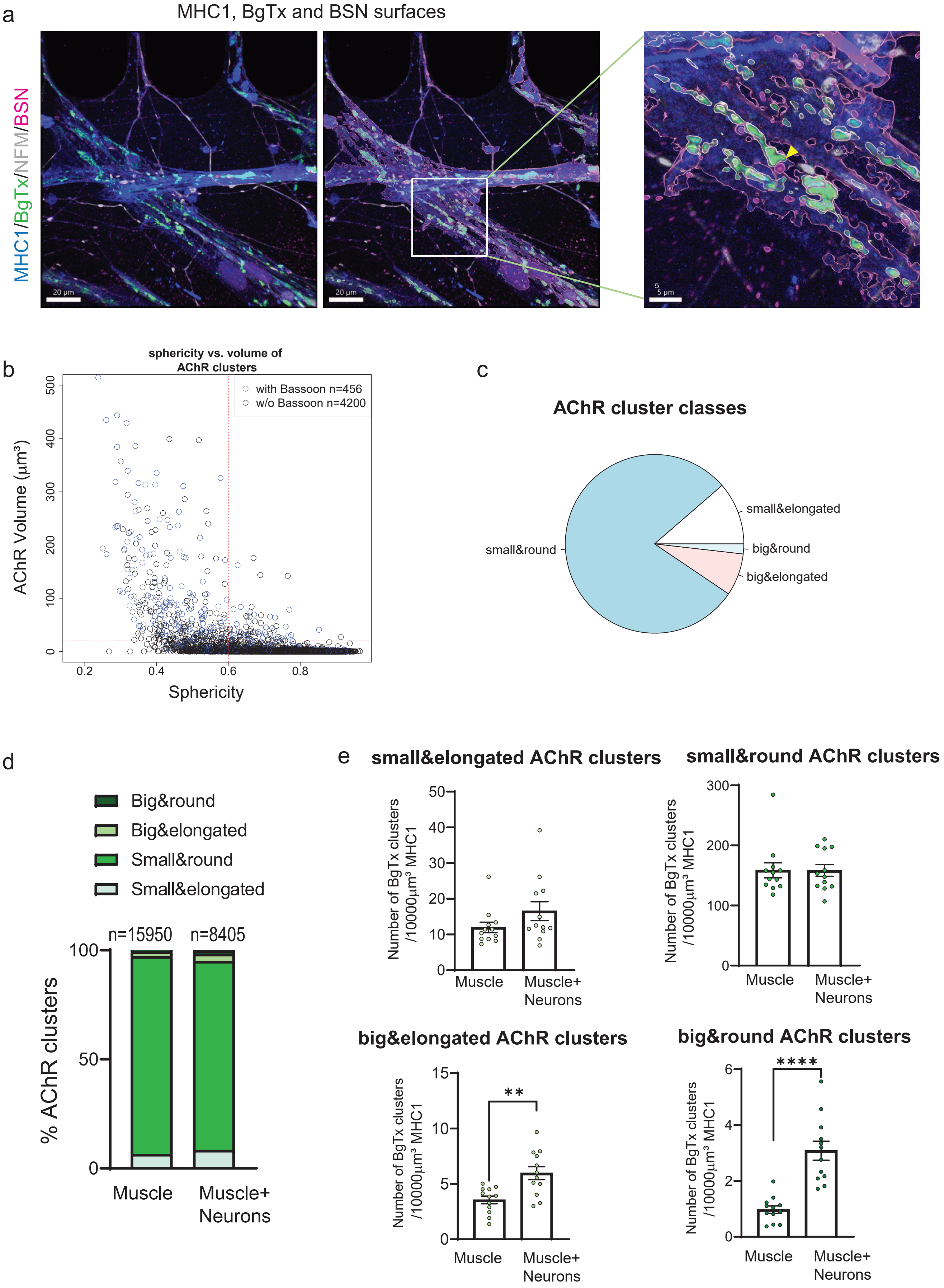

a
MHC1, BgTx and BSN surfaces
MHC1/BgTx/NFM/BSN
20 µm
20 µm
5 µm
b
sphericity vs. volume of AChR clusters
with Bassoon n=456
w/o Bassoon n=4200
AChR Volume (µm³)
Sphericity
c
AChR cluster classes
small&elongated
big&round
big&elongated
small&round
d
Big&round
Big&elongated
Small&round
Small&elongated
n=15950
n=8405
% AChR clusters
Muscle
Muscle+ Neurons
e
small&elongated AChR clusters
small&round AChR clusters
big&elongated AChR clusters
big&round AChR clusters
Number of BgTx clusters /10000µm³ MHC1
Muscle
Muscle+ Neurons
**
****

Suppl. Figure 5

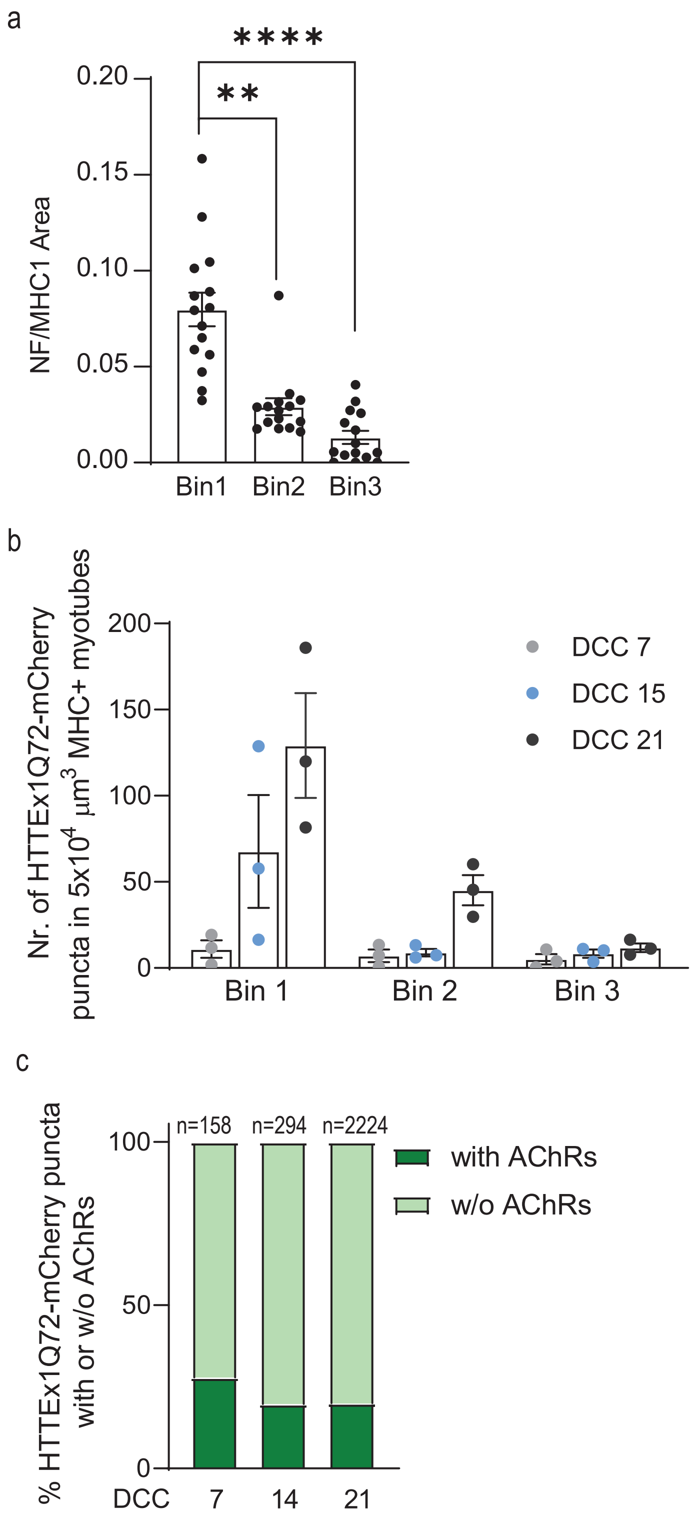
